# A natural mutator allele shapes mutation spectrum variation in mice

**DOI:** 10.1101/2021.03.12.435196

**Authors:** Thomas A. Sasani, David G. Ashbrook, Annabel C. Beichman, Lu Lu, Abraham A. Palmer, Robert W. Williams, Jonathan K. Pritchard, Kelley Harris

## Abstract

Although germline mutation rates and spectra can vary within and between species, genetic modifiers of these traits have long eluded detection. In this study, we searched for loci that influence germline mutagenesis using a uniquely powerful resource: a panel of recombinant inbred mouse lines known as the <u>B</u>X<u>D</u>, descended from the laboratory mouse strains C57BL/6J (<u>B</u>) and DBA/2J (<u>D</u>). Each BXD lineage has been maintained by brother-sister mating in the near absence of natural selection, accumulating *de novo* mutations for up to 50 years on a known genetic background that is a unique linear mosaic of *B* and *D* haplotypes. We show that mice inheriting *D* haplotypes at a quantitative trait locus (QTL) on chromosome 4 accumulate C>A germline mutations at a 50% higher rate than those inheriting *B* haplotypes, primarily due to the activity of a C>A-dominated mutational signature known as SBS18. The *B* and *D* QTL haplotypes encode different alleles of the DNA repair gene *Mutyh*, which underlies the heritable colorectal cancer syndrome in which SBS18 was first identified. The *B* and *D Mutyh* alleles are present in wild populations of *Mus musculus domesticus*, providing evidence that common genetic variation modulates germline mutagenesis in a model mammalian species.

## Introduction

Although all living organisms maintain low mutation rates via conserved DNA repair and proofreading pathways, the fidelity of genetic inheritance varies by orders of magnitude across the tree of life ^1^. Evolutionary biologists have long debated why mutation rates vary so dramatically, citing trade-offs including the necessity of beneficial mutations for adaptation ^2^, the cost of DNA replication fidelity ^3^, and the inefficiency of selection against weak mutation rate modifiers ^1^.

Within humans, germline mutation rates also vary among families ^4–6^ and are particularly elevated in individuals affected by a rare heritable cancer syndrome ^7^. Human populations also exhibit variation in the *mutation spectrum* ^*8,9*^, a summary of the relative abundances of specific base substitution types (C>A, C>T, A>G, etc.) ^8–10^. Genetic mutation rate modifiers, or “mutator alleles”, have been invoked as possible contributors to these patterns; however, the relative importance of genetic and environmental mutators remains poorly understood ^6,11^.

Previous attempts to study the genetic architecture of germline mutation rates have been hindered in part by the dependence of mutation rates on parental age ^4,12,13^. Our study avoids this and other confounders by analyzing a large family of recombinant inbred mouse lines (RILs), whose environments and generation times have been controlled by breeders for decades. Beginning in 1971, crosses of two inbred laboratory mouse lines—C57<u>B</u>L/6J and <u>D</u>BA/2J—were used to generate several cohorts of B-by-D (BXD) recombinant inbred progeny ^14^. These progeny have accumulated *de novo* mutations during many generations of sibling inbreeding, much like the members of mutation accumulation (MA) lines commonly used to measure mutation rates in microorganisms and invertebrates ^15^.

## Results

### Identifying germline mutations in BXDs

The BXD family was generated via six breeding epochs initiated between 1971 and 2014 ^14^ (**Extended Data Figure 1)**; each epoch contains between 7 and 49 recombinant inbred lines (RILs). We sequenced the genome of a whole spleen from each BXD RIL, excluded lines confounded by significant heterozygosity (including all of epoch 6), and retained 94 lines that had each been inbred for at least 20 generations (**Extended Data Table 1, Extended Data Fig. 2, Supplementary Information**). We identified 63,914 single nucleotide variants (SNVs) that were homozygous for a non-reference allele in one RIL and homozygous for the reference allele in the C57BL/6J and DBA/2J parents, as well as all other BXDs; each autosomal “singleton” likely arose as a *de novo* germline mutation during inbreeding of the RIL in which it appears (**Figure 1a**). Across BXD lines, singleton counts are positively correlated with the number of generations of inbreeding (Poisson regression *p* < 2.2 × 10^-16^, **Fig. 1a**). As reported in other inbred mice ^16^, the high density of singletons in conserved genomic regions suggests that the effects of purifying selection have been minimal during BXD inbreeding (Kolmogorov-Smirnov test, *p* < 2.2 × 10^-16^) **(Supplementary Information, Extended Data Fig. 3**).

**Figure 1:**
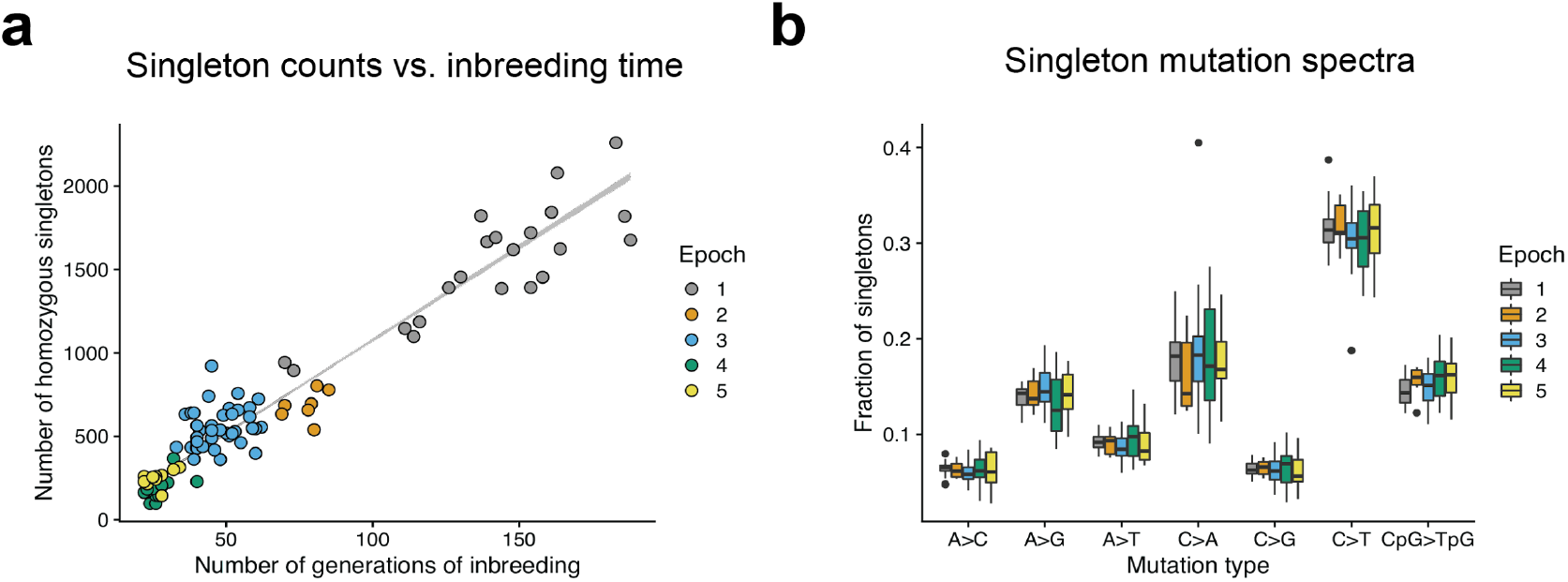
Accumulation of homozygous singletons over many generations of laboratory inbreeding. (a) Counts of autosomal homozygous singletons in 94 BXDs correlate with the number of generations of inbreeding. Lower-numbered epochs are older and have been inbred for more generations. Line is from a Poisson regression (identity link) with 95% confidence bands. (b) Fractions of singletons from each epoch that belong to each of seven mutation types, including the six possible transitions and transversions as well as CpG>TpG. Strand complements are collapsed. Boxes represent the median and interquartile range (IQR), and whiskers extend to 1.5 times the IQR. The strain with an extremely high fraction of C>A singletons is BXD68.

### A QTL for the C>A mutation rate

Mutation spectra inferred from BXD singletons are similar to spectra previously inferred from *de novo* germline mutations in mice^17^, but we observed variation within epochs (**Supplementary Information, Fig. 1b**). We hypothesized that some of this variation might be caused by mutator loci, where *B* and *D* alleles have different functional impacts on DNA repair or replication fidelity. To test this hypothesis, we performed quantitative trait locus (QTL) mapping using R/qtl2^18^ for the overall mutation rate in each line, and for the rates and fractions of the seven mutation types shown in **Fig. 1b** (**Supplementary Table 1**). We excluded BXD68 from our QTL scans due to its exceptional C>A singleton rate and fraction (**Fig. 1b**).

We did not find any genome-wide significant QTL for the overall mutation rate (**Extended Data Fig. 4a**), but a scan for loci associated with the fraction of C>A singleton mutations revealed a highly significant peak on chromosome 4 (**Fig. 2a**; maximum LOD of 17.9 at 116.8 Mbp; Bayes 95% C.I. = 114.8 –118.3 Mbp). BXD lines with *D* haplotypes at this locus (hereafter called *D* lines; *n* = 56) have substantially more C>A mutations than lines with *B* haplotypes (hereafter *B* lines; *n* = 38) (**Fig. 2b;** *p* < 2.2 × 10^-16^), an effect that explains 59.2% of the variance in BXD C>A singleton fractions. We discovered the same LOD peak via a QTL scan for the C>A mutation rate (**Fig. 2a**; maximum LOD of 6.9 at 116.8 Mbp; Bayes 95% C.I. = 114.8 –118.8 Mbp). On average, the *D* lines have accumulated C>A mutations at a rate of 1.22 × 10^-9^ per base pair per generation (95% CI: 1.08 - 1.37 × 10^-9^), over 1.5-fold higher than the rate of 7.32 × 10^-10^ (95% CI: 6.66 - 8.11 × 10^-10^) observed in *B* lines. This C>A rate difference gives the *D* lines a 1.11-fold higher overall mutation rate than the *B* lines, but is not large enough to produce a globally significant association between the C>A QTL and the overall mutation rate. No other mutagenesis-related QTL scans identified genome-wide significant peaks (**Extended Data Fig. 4b**). In a principal component analysis, variation in C>A fractions largely drives PC1, which separates the *B* lines from the *D* lines (**Fig. 2c**). Since a higher C>A fraction distinguished the DBA/2J and C57BL/6NJ mutation spectra in a previous report^16^, the observed QTL on chromosome 4 appears to fit the profile of a mutator locus responsible for a major difference between the parental strains’ mutation spectra.

**Figure 2:**
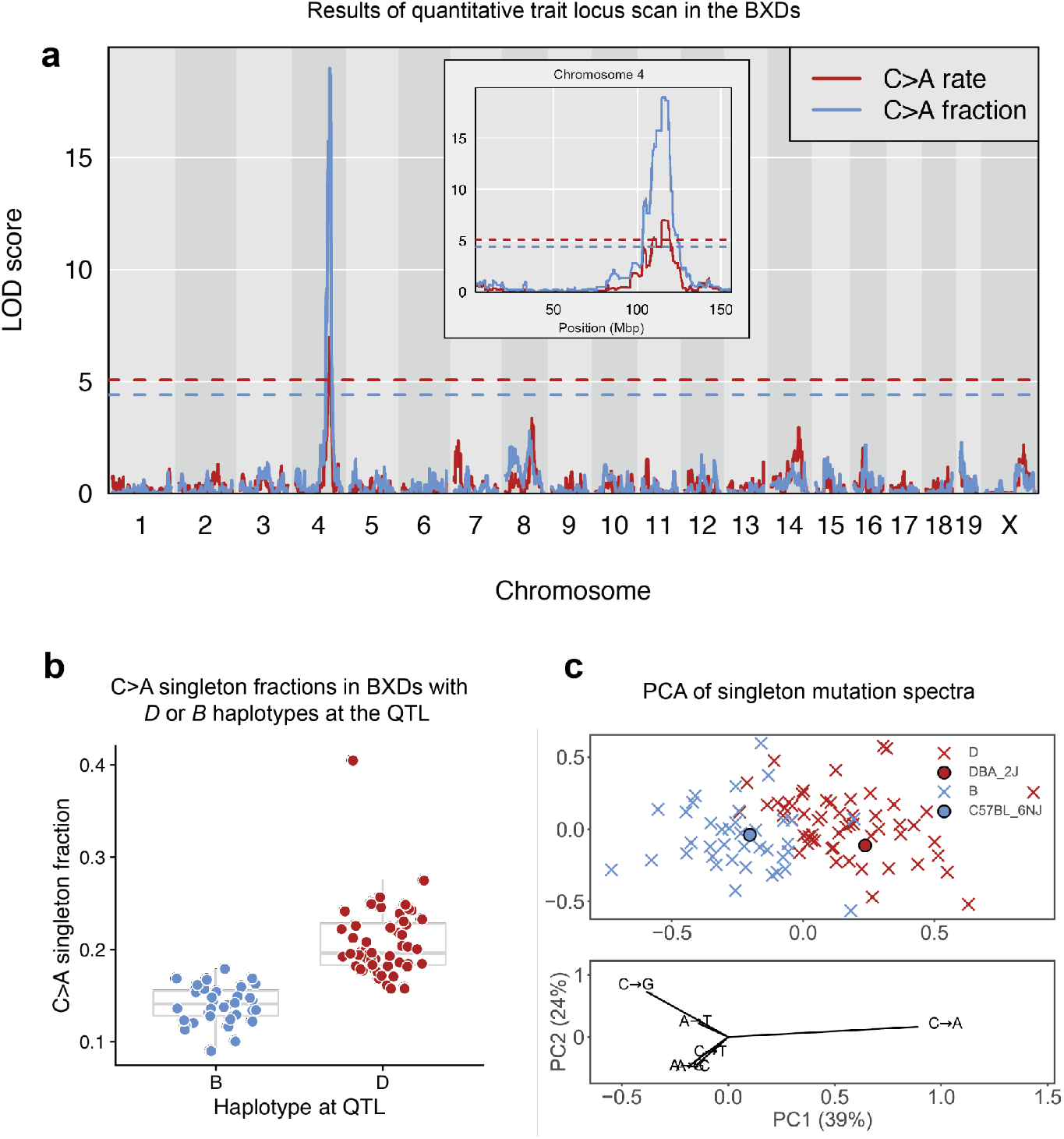
A quantitative trait locus on chromosome 4 for the germline C>A mutation rate. (a) LOD scores for the centered log-ratio transformed fraction (blue) or estimated rate (red) of C>A mutations. Blue and red dashed lines indicate genome-wide significance thresholds (using 1,000 permutations and a Bonferroni-corrected α = 0.05/15) for the fraction and rate scans, respectively. C>A fraction and rate phenotypes are included in the GeneNetwork database as BXD_24430 and BXD_24437, respectively. Inset is zoomed to only show LOD scores on chromosome 4. (b) C>A singleton fractions in BXD lines homozygous for the *D* (*n* = 56) or the *B* haplotype (*n* = 38) at the QTL on chromosome 4. Boxes represent the median and interquartile range (IQR), and whiskers extend to 1.5 times the IQR. C>A fraction outlier BXD68 was not included in the QTL scan. (c) Principal component analysis of the 6-dimensional mutation spectra of BXD singletons (*n* = 94 lines). BXDs are denoted as crosses colored by parental ancestry at the QTL on chromosome 4 (*D* [red] or *B* [blue]). Strain-private mutation spectra for DBA/2J and C57BL/6NJ^16^ are denoted with circles. Loadings are plotted for each mutation type below. Fractions of each mutation type were centered log-ratio transformed prior to PCA.

### The QTL contains coding variation in *Mutyh*

Using SnpEff ^19^, a tool that predicts the impact of genetic variation on protein function, we identified 61 MODERATE-impact and 5 HIGH-impact sequence differences between *B* and *D* haplotypes within the QTL, affecting 21 out of 76 protein-coding genes in the locus (**Supplementary Information**). Only one of these 21 genes is annotated by the Gene Ontology resource ^20,21^ as being relevant to “DNA repair” or the “cellular response to DNA damage”: the mouse homolog of the mutY DNA glycosylase, *Mutyh. Mutyh* is required for base excision repair of 8-oxoguanine (8-oxoG) lesions, which are a result of damage to guanine nucleotides by reactive oxygen species. Left unrepaired, 8-oxoG frequently base pairs with adenine rather than thymine, causing C>A mutations during subsequent DNA replication ^22^. We observed a total of 5 MODERATE-impact differences between the *B* and *D* alleles of *Mutyh* (**Table 1**).

**Table 1:** *Mutyh* missense mutations in the BXD family. Amino acid positions are relative to Ensembl transcript ENSMUST00000102699.7.

| Amino acid change | Coordinates in mm10 | Fixed on *D* haplotypes? |
| --- | --- | --- |
| p.Gln5Arg | Chr4:116814338 | Yes |
| p.Arg24Cys | Chr4:116814394 | Yes |
| p.Ser69Arg | Chr4:116815658 | Yes |
| p.Arg153Gln | Chr4:116816476 | No (private to BXD68) |
| p.Thr312Pro | Chr4:116817416 | Yes |
| p.Ser313Pro | Chr4:116817419 | Yes |

*Mutyh* deficiency contributes to a mutator phenotype in the germlines of TOY-KO mice, a triple knockout strain lacking *Mutyh* as well as *Mth1* and *Ogg1*, the other primary genes involved in genomic 8-oxoguanine repair ^23^. TOY-KO mice have a *de novo* germline mutation rate elevated nearly 40-fold above normal ^23^ and a *de novo* mutation spectrum with very high cosine similarity (0.94) to SBS18, a mutational signature dominated by CA>AA and CT>AT mutations ^24,25^ that was first identified in colorectal and pancreatic tumors from human patients with pathogenic germline *MUTYH* mutations ^24,26,27^.

We used SigProfilerExtractor^28^ to decompose the BXD singleton mutation spectra into combinations of COSMIC mutation signatures previously discovered in human cancers and found that 13.9% of BXD singletons were assigned to the *MUTYH-*associated SBS18 signature (**Supplementary Table 2**). SigProfilerExtractor decomposed the remaining singletons into three additional signatures: SBS1 (14.2% of mutations), SBS5 (55.6%), and SBS30 (16.4%). SBS1, SBS5, and SBS30 were each identified in the majority of BXDs (94/94, 91/94, and 78/94 BXDs, respectively), with no statistically significant imbalances between *B* and *D* lines (all Chi-square *p* values = 1.0). In contrast, SBS18 was identified in just 52/94 lines, including 50/56 *D* haplotype lines and only 2/38 *B* haplotype lines (Chi-square *p* = 4.9 *x* 10^-15^). SBS18 activity is thus a highly accurate classifier of BXD haplotype status at the QTL on chromosome 4, and as expected, *D* lines are enriched for the same 3-mer C>A mutation types that are most abundant in TOY-KO germline mutations and the SBS18 signature (**Fig. 3**).

**Figure 3:**
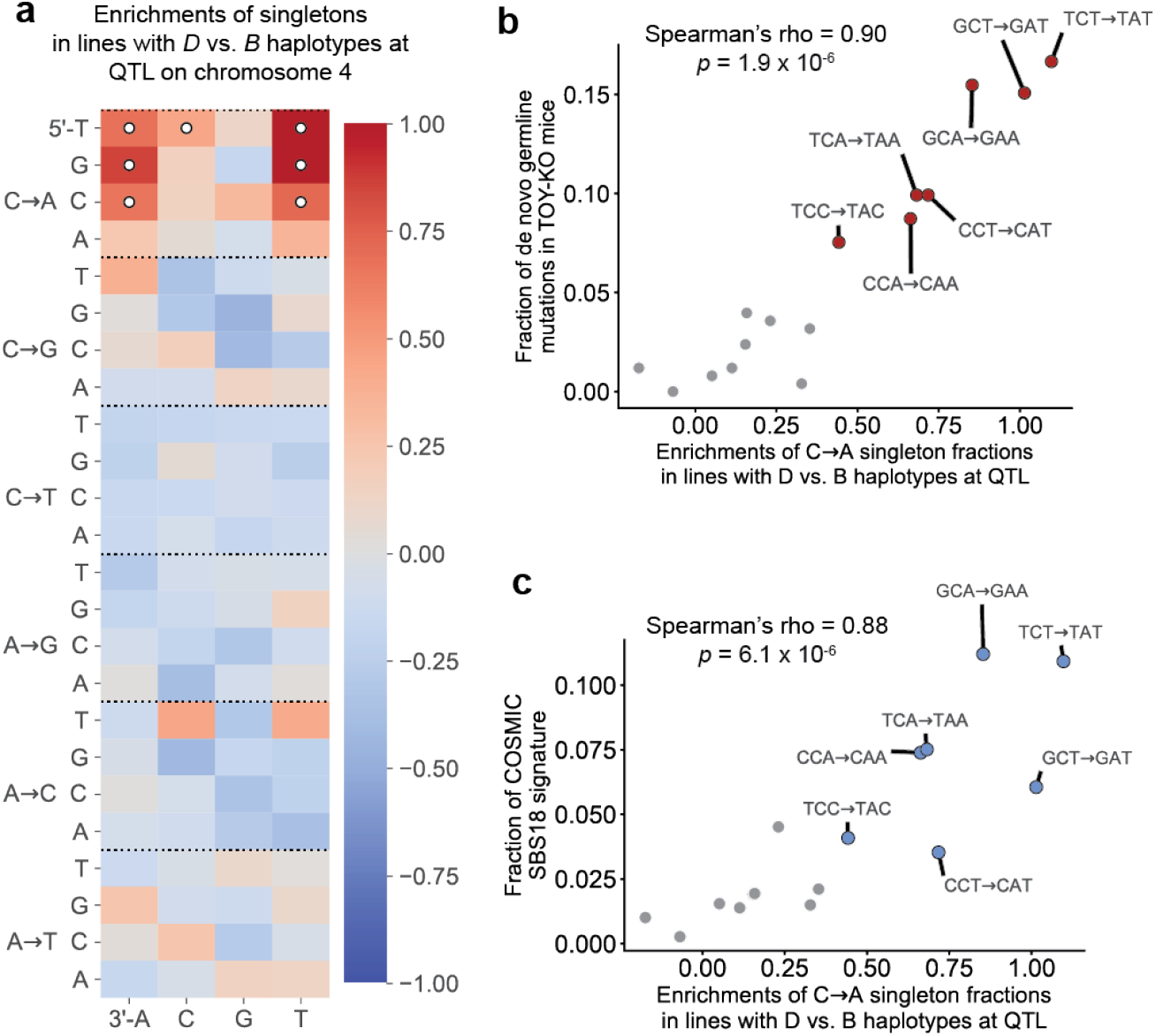
Particular C>A mutation types are more frequent in lines with the *D* haplotype at the C>A QTL. (a) Log-2 ratios of mutation fractions in lines with *D* haplotypes compared to lines with *B* haplotypes at the QTL on chromosome 4. Mutation types with Chi-square test of independence *p*-values < 0.05/96 (Bonferroni-corrected) are marked with white circles. (b) The log-2 enrichments of C>A mutations in each 3-mer context in *D* vs. *B* lines from heatmap (a) are plotted against the relative abundances of C>A mutations in each context in a previously reported set of *de novo* mutations from *Mutyh*^*-/-*^*/Ogg1*^*-/-*^*/Mth1*^*-/-*^ mice^23^ (*n* = 252 mutations). 3-mer mutation types with significant enrichments labeled in (a) are colored in red and outlined in black. (c) The log-2 enrichments of C>A mutations in each 3-mer context in *D* vs. *B* lines from heatmap (a) are plotted against the relative abundances of C>A mutations in each context in the SBS18 COSMIC mutation signature. 3-mer mutation types with significant enrichments labeled in (a) are colored in blue and outlined in black.

### Alternative explanations for the C>A QTL

We note that three other genes within the QTL interval are associated in the Gene Ontology database with “DNA repair” or the “cellular response to DNA damage” (*Plk3, Rad54L*, and *Dmap1*). One additional gene, *Prdx1*, is involved in the “cellular response to oxidative stress”. However, none of these genes harbors nonsynonymous coding differences between *D* and *B* haplotypes, and none to our knowledge is implicated in C>A mutagenesis, making them *a priori* less likely than *Mutyh* to cause the observed mutator phenotype.

We also used GeneNetwork ^29^ to test for associations between the SNP marker with the highest LOD score at the QTL (rs52263933) and the expression of genes that might contribute to the C>A mutator phenotype (**Supplementary Information**). We queried gene expression in a number of cell types, including kidney, gastrointestinal tissue, and hematopoietic stem cells; in these tissues, the most significant correlations were between rs52263933 genotypes and expression of *Atpaf1, Rps8*, and *Mutyh*, respectively. Although expression quantitative trait loci (eQTLs) may contribute to the C>A germline mutator phenotype, our analysis is limited by a number of factors: namely, that our QTL scan implicated a relatively large (approximately 4 Mbp) haplotype segment, and that we are unable to query BXD expression data from germline tissues such as testis or ovary.

Finally, we queried the database of structural variants (SVs) identified by the Mouse Genomes Project consortium ^30^ but did not find any fixed structural differences between *B* and *D* haplotypes within the QTL interval that might explain the C>A mutator phenotype (**Supplementary Information**).

### A C>A hypermutator phenotype in BXD68

One outlier *D* line, BXD68, was excluded from QTL scans because its singleton C>A fraction was 5.6 standard deviations above the mean (**Fig. 2b)**. SigProfilerExtractor assigned nearly 55% of BXD68’s mutations to SBS18, suggesting a shared etiology between its hypermutator phenotype and the mutator phenotype common to all *D* strains. We hypothesized that BXD68 might harbor a private mutator allele within the chromosome 4 QTL, and discovered two BXD68 singletons within this interval: an intronic variant in *Kdm4a* and, strikingly, a missense mutation in *Mutyh* (p.Arg153Gln) (**Table 1**). One DNA repair gene outside the QTL, *Rev3l*, harbors a nonsynonymous singleton in BXD68, but it is located on chromosome 10 and is associated with a mutational signature dominated by mutations at GC dinucleotides ^31^ that does not resemble any mutator phenotype active in the BXD lines.

The BXD68 singleton affects an amino acid that is conserved between humans and mice (p.Arg179, relative to the human Ensembl transcript ENST00000372098.3). Two missense mutations that affect the human p.Arg179 amino acid (rs747993448 and rs143353451) are both listed in the ClinVar database as being pathogenic or likely pathogenic^32^, and the murine p.Arg153Gln amino acid change is predicted to be deleterious by both PROVEAN^33^ and SIFT^34^. Based on this evidence, we hypothesize that p.Arg153Gln arose as a *de novo* germline mutation in BXD68 and impairs the 8-oxoguanine DNA damage response even more severely than the mutator allele(s) that occurs on the *D* haplotype.

### Both *B* and *D Mutyh* alleles segregate in wild mice

Using publicly available whole genome variation data, we found all five nonsynonymous differences between the *B* and *D Mutyh* alleles to be segregating in wild populations of *Mus musculus domesticus*, the subspecies from which laboratory mice derive most of their genetic ancestry^35^ (**Fig. 4a**), This suggests that the *D* haplotype mutator may be shaping the accumulation of genetic variation in nature (though no wild mice are known to possess the BXD68-private p.Arg153Gln variant). Unexpectedly, the outgroup species *Mus spretus* appears to be fixed for the *D* allele at four of the five missense sites in *Mutyh* (**Fig. 4a**). The *D* mutator allele therefore appears more likely to be ancestral than the reference *B* allele, a conclusion supported by a multiple sequence alignment of additional vertebrates (**Fig. 4b**).

**Figure 4:**
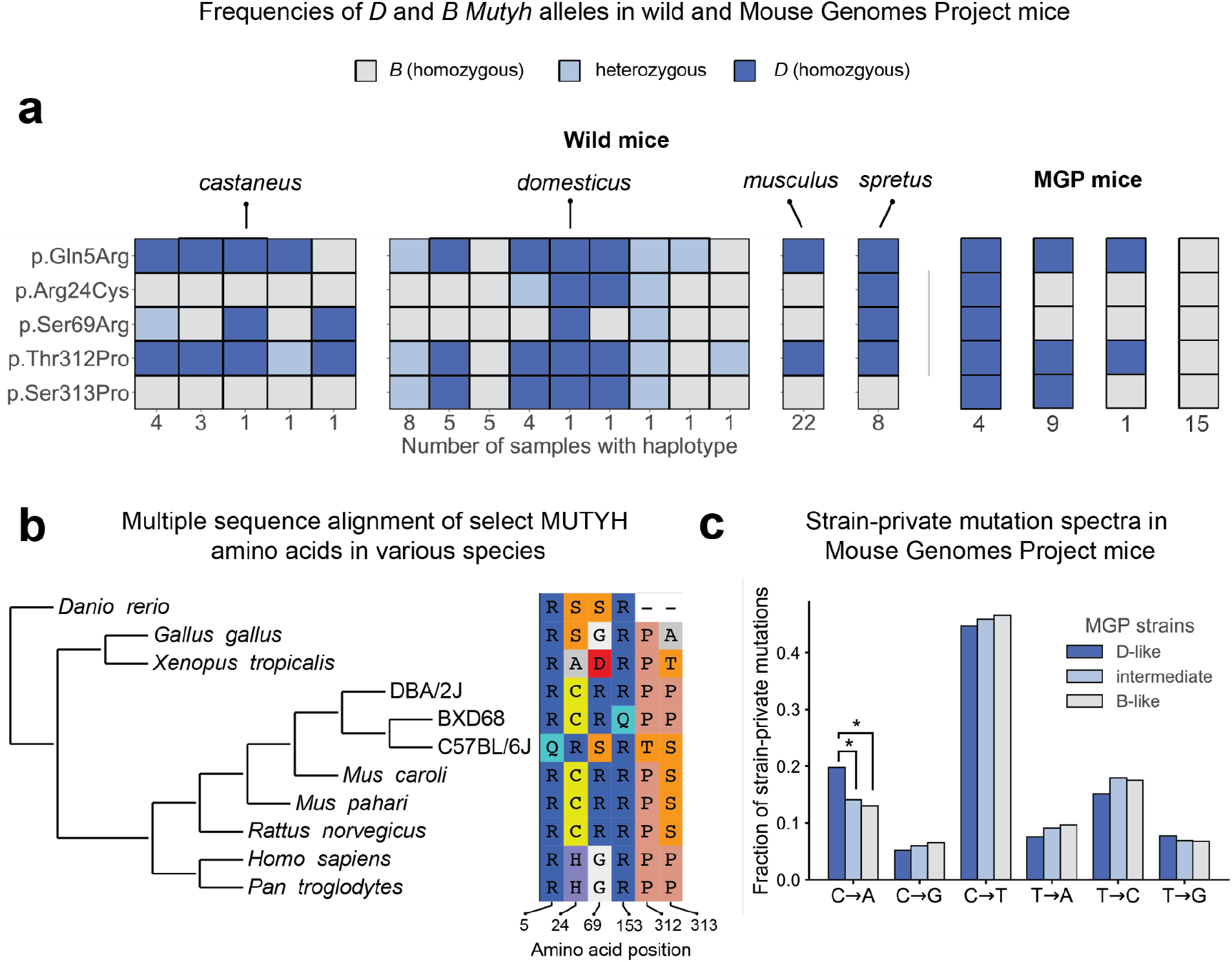
Nonsynonymous differences between *B* and *D Mutyh* alleles segregate in both wild and inbred mouse strains and appear to be ancestral in DBA/2J. (a) Presence of *D* or *B Mutyh* alleles in 67 wild mice^41^ and in Sanger Mouse Genomes Project (MGP) strains that have associated strain-private singleton data^16^. Unique combinations of *Mutyh* alleles were identified in each wild or MGP mouse, and are represented using columns of colored boxes. Wild mice are grouped by species or subspecies, and the number of mice with each unique combination of *Mutyh* alleles is listed below each column. MGP mice are grouped by unique combinations of *Mutyh* alleles, and the number of mice with each combination is listed below each column. (b) Multiple sequence alignment of MUTYH amino acids is subsetted to only show the six amino acids affected by MODERATE or HIGH-impact mutations in the BXD. Positions of amino acids in the murine MUTYH peptide sequence (ENSMUST00000102699.7) are shown below each column. (c) Comparison of strain-private mutation spectra in MGP strains^16^ with *D*-like, intermediate, or *B*-like genotypes at *Mutyh* missense mutations. *P*-value of Chi-square test comparing C>A counts between *D*-like and intermediate strains: 3.3 × 10^-7^; between *D*-like and *B*-like strains: 1.4 × 10^-10^; between intermediate and *B*-like strains: 1.2 × 10^-1^.

Among the 29 laboratory mouse strains sequenced by the Sanger Mouse Genomes Project (MGP)^30^, four (including DBA/2J) match the *D* lines at all five sites, while 15 (including C57BL/6NJ) match *B* lines at all five sites. Nine “intermediate” strains harbor *D* alleles at amino acids 5, 312, and 313 but harbor *B* alleles at amino acids 24 and 69 (**Fig. 4a**). As expected, the *D-*like strains have the highest fractions of recent germline C>A mutations, particularly of the CA>AA and CT>AT dinucleotide types that are enriched in SBS18 (**Fig. 4c, Extended Data Fig. 5**). However, we found no significant mutation spectrum differences between the intermediate and *B*-like strains (**Fig. 4c**). These observations tentatively point to p.Arg24Cys and p.Ser69Arg as the variants most likely to underlie the QTL mutator phenotype.

In theory, *B* and *D* alleles should shape natural mouse genetic variation by causing more C>A variants to accumulate in wild populations with more *D* alleles. Although we found some evidence for elevated C>A mutagenesis in wild mouse subspecies with the highest *D* allele frequencies (**Supplementary Information, Extended Data Fig. 6-8**), other forces such as biased gene conversion and additional mutators might contribute to this pattern. Additional sampling of wild mice will be needed to better assess the historical activity of the BXD mutator.

### No evidence for selection on *Mutyh* alleles

Since new mutations are more often deleterious than beneficial, natural selection is generally expected to favor lower mutation rates ^1^. However, we found no evidence for deviations from neutrality near the QTL in wild mice (**Supplementary Information**). This finding is somewhat surprising in light of the mutator’s effect size; using population genetic theory and a previous estimate of the average fitness effect of *de novo* coding mutations in mice^36^, we estimated that the *B* allele should avoid enough excess deleterious mutations to be favored with a selection coefficient (*s*) of about 3 × 10^-4^ to 6 × 10^-4^, as long as it is semi-or fully dominant over the *D* allele (**Supplementary Information**). In theory, this should be advantageous enough to sweep to fixation in a mouse population of effective size *N* = 5 × 10^4 37,38^. However, a number of factors may have impeded such a sweep, including mouse population substructure, the activities of other genetic mutation rate modifiers, and antagonistic pleiotropy. Additionally, if the *D* allele’s mutagenic effect is recessive (like the disease phenotypes associated with deleterious human *MUTYH* missense mutations), the ancestral mutator may hide out neutrally in heterozygotes, impeding fixation of the derived *B* allele.

## Discussion

Our discovery of a genetic modifier of the murine C>A mutation rate provides new support for the long-standing theoretical prediction that multicellular eukaryotes have a limited ability to optimize their germline replication fidelity ^1^. Our work shows that common mutator alleles, previously identified only in microorganisms such as *Saccharomyces cerevisiae* ^*39,40*^, also shape vertebrate genetic diversity.

We argue that the BXD mutator phenotype is likely caused by natural variation in *Mutyh*, the only DNA repair gene in the QTL interval that contains nonsynonymous coding differences between the parental strains. Although we cannot completely rule out the contributions of nearby genes or regulatory variants, *Mutyh* exhibits the strongest prior link to the C>A dominated SBS18 mutation signature of any protein-coding gene in the QTL interval (**Supplementary Information**). In human patients with colorectal cancer, SBS18 activity has been found to be 100% predictive of inherited pathogenic biallelic *MUTYH* missense variants ^26^. *Mutyh* is also the only gene in the QTL interval (and one of only two DNA repair genes genome-wide) that harbors non-synonymous coding variation in BXD68, an outlier line with an exceptionally high C>A mutation rate and SBS18 burden. Other than *Mutyh*, none of the other genes within the C>A QTL has a documented association with SBS18, and none would parsimoniously explain the C>A hypermutator phenotype of BXD68 (**Supplementary Information**).

Our findings add weight to the conjecture that natural mutator alleles underlie some of the species-specific and population-specific signatures previously observed in humans and other great apes ^9,10^, and demonstrate that mutators are mappable in model organisms using QTL analysis. Differences in mutation spectra observed across other mouse populations ^16^ suggest that the BXD mutator is just one of several active mutator alleles in mice, any of which might have been detected if the “right” parents had been selected to initiate a cross like the BXD. We anticipate that mutator allele discovery will become increasingly feasible across the tree of life as sequencing costs continue to decline, providing long-awaited data needed to test theoretical predictions about selection on this fundamental phenotype.

## Online Methods

All Python and R code used in these analyses is available at https://github.com/tomsasani/bxd_mutator_manuscript. We used *snakemake* ^*1*^ to write a collection of pipelines that can be used to reproduce all analyses described in the manuscript.

### Construction of the BXD RILs

Detailed descriptions of the BXD dataset, including the construction of the BXD recombinant inbred lines, can be found in a previous manuscript ^2^. Briefly, the BXD RILs were derived from crosses of the DBA/2J and C57BL/6J inbred laboratory strains initiated in six distinct “epochs” from 1971 to 2014. RILs were produced using one of two strategies: four epochs were produced using the standard F2 cross, and two were produced using the advanced intercross strategy. In the F2 cross design, a male DBA/2J mouse is crossed to a C57BL/6J female to produce F1 animals that are heterozygous for parental ancestry at essentially all loci in the genome. Pairs of these F1 animals are then crossed to produce F2s. To generate each individual recombinant inbred line, a brother and sister are picked from among the F2s and mated; this brother-sister mating strategy continues for many generations. In the AIL cross design, F2s are generated as in the standard F2 cross. However, pseudo-random pairs of F2 animals are then crossed to generate F3s, pseudo-random pairs of F3s are crossed to generate F4s, and so on, for up to 14 generations. Then, to generate inbred lines, brother-sister matings are once again initiated from the offspring of the final pseudo-random cross. Schematic diagrams of the F2 cross and AIL strategies are shown in **Extended Data Figure 1**.

### Whole-genome sequencing, alignment, and variant calling

BXD mice (all males) were euthanized using isoflurane. Spleen tissue was collected immediately, flash frozen with liquid nitrogen, and placed in a -80 degree freezer for subsequent analysis. All DNA extraction, library preparations and sequencing was carried out by HudsonAlpha (Huntsville, AL, USA). High molecular weight genomic DNA was isolated from 50 to 80 mg of spleen tissue using the Qiagen MagAttract kit (Qiagen, Germantown, MD). The Chromium Gel Bead and Library Kit (v2 HT kit, revision A; 10X Genomics, Pleasanton, CA, USA) and the Chromium instrument (10X Genomics) were used to prepare libraries for sequencing; barcoded libraries were then sequenced on the Illumina HiSeq X10 system. FASTQ files were aligned to the mm10/GRCm38 reference genome using the 10X LongRanger software (v2.1.6), using the “Lariat” alignment approach. Variant calling was carried out on aligned BAM files using GATK version v3.8-1-0-gf15c1c3ef ^3^ to generate gVCF files; these gVCFs were then joint-called to produce a complete VCF file containing variant calls for all BXDs and founders. GATK Variant Quality Score Recalibration (VQSR) was then applied to the joint-called VCF. A list of known, “true-positive” variants was created for VQSR by identifying variants which were shared across three distinct call sets: 1) variants identified in DBA/2J in this study 2) variants previously identified in DBA/2J^4^ and 3) variants identified in DBA/2J in the Sanger Mouse Genomes Project ^5^. This generated a set of 3,972,727 SNPs, 404,349 deletions and 365,435 insertions; we were highly confident that these varied between the DBA/2J and reference sequences and expected that each should appear in approximately 50% of the BXD strains. The SNP and indel variant calls from the Sanger Mouse Genomes project ^5^ were also used as a training resource for VQSR.

### Identifying homozygous singleton variants in the BXD RILs

To confidently identify singletons (sites with a non-reference allele in exactly one of the BXD recombinant inbred lines) we iterated over all autosomal variants in the joint-genotyped VCF using *cyvcf2* ^6^ and identified variants that passed the following filters: first, we removed all variants that overlapped segmental duplications or simple repeat annotations in mm10/GRCm38, which were downloaded from the UCSC Genome Browser. We limited our analysis to single nucleotide variation and did not include any small insertion or deletion variants. At each site, we required both founder genotypes (DBA/2J and C57BL/6J) to be homozygous for the reference allele, for each of these genotypes to be supported by at least 10 sequencing reads, and for Phred-scaled genotype qualities in both founders to be at least 20. We then required that exactly one of the BXD RILs had a heterozygous or homozygous alternate (i.e., non-reference) genotype at the site; although we only included 94 BXDs in downstream analyses (**Supplementary Information**), the genome sequences of all sequenced BXDs (except for the small number of BXDs that were isogenic with another line) were considered when identifying potential singletons to ensure that none of the excluded strains possessed a putative singleton allele present in the focal strain. To include a heterozygous genotype in our singleton callset, we required its allele balance (the fraction of reads supporting the non-reference allele) to be >= 0.9. For candidate heterozygous and homozygous singletons, we also required the genotype call to be supported by at least 10 total sequencing reads (including both reference and alternate alleles) and have Phred-scaled genotype quality at least

20. Finally, we confirmed that at least one other BXD shared a parental haplotype identical-by-descent with the focal strain (i.e., the strain with the putative singleton) at the singleton site but was homozygous for the reference allele at that site (**Supplementary Information**).

We additionally annotated the full autosomal BXD VCF with SnpEff ^7^ version 4.3t, using the GRCm38.86 database and the following command:

~~~
java -Xmx16g -jar /path/to/snpeff/jarfile GRCm38.86 /path/to/bxd/vcf > /path/to/uncompressed/output/vcf
~~~

### Annotating singletons with triplet sequence contexts and conservation scores

For each candidate singleton variant, we were interested in characterizing the 5’ and 3’ sequence context of the mutation, as well as the phastCons conservation score of the nucleotide at which the variant occurred. To determine the sequence context of each variant, we used the *mutyper* python API ^8^. To annotate each variant with its phastCons score, we downloaded phastCons scores derived from a 60-way placental mammal alignment for the mm10/GRCm38 genome build in WIG format from the UCSC Table Browser. We then converted the WIG files to BED format using the *bedops wig2bed* subcommand ^*9*^, compressed the BED format files with bgzip, and indexed the compressed BED files with tabix. Within the Python script used to identify singletons, we then used *pytabix* (https://github.com/slowkow/pytabix) to query the phastCons BED files at each putative singleton.

### Quantitative trait locus mapping

We used the R/qtl2 software ^10^ for QTL mapping in this study. Prior to running QTL scans, we downloaded a number of data files from the R/qtl2 data repository (https://github.com/rqtl/qtl2data), including physical (Mbp) and genetic (cM) maps of the 7,320 genotype markers used for QTL mapping, as well as a file containing genotypes for all BXDs at each of these markers. These files are also included in the GitHub repository associated with this manuscript.

We inserted pseudo-markers into the genetic map using *insert_pseudomarkers* and calculated genotype probabilities at each marker using *calc_genoprob*, with an expected error probability of 0.002. We additionally constructed a kinship matrix describing the relatedness of all strains used for QTL mapping using the leave-one-chromosome-out (LOCO) method. We then performed a genome scan using a linear mixed model (*scan1* in R/qtl2), including the kinship matrix, a covariate for the X chromosome, and two additive covariates. The first additive covariate denoted the number of generations each RIL was intercrossed prior to inbreeding (0 for strains derived from standard F2 crosses, and *N* for strains derived from advanced intercross, where *N* is the number of generations of pseudo-random crosses performed before the start of inbreeding), and the second additive covariate denoted the epoch from which the strain was derived. To assess the significance of any log-odds peaks, we performed a permutation test (1,000 permutations) with *scan1perm*, using the same covariates and kinship matrix as described above. We calculated the Bayes 95% credible intervals of all peaks using the *bayes_int* function, with *prob=0*.*95*.

When performing QTL scans for a particular mutation fraction, we treated the phenotype as the centered log-ratio transform of the fraction of singletons of that type in each strain. When performing scans for mutation rates, we used the untransformed mutation rates (per base pair per generation) as the phenotype values.

### Comparing C>A singleton fractions between BXDs with *D* and *B* haplotypes at the QTL on chromosome 4

To compare singleton fractions in BXDs with either *D* or *B* haplotypes at the QTL on chromosome 4, we first used a simple Welch’s two-sided t-test, which returned *p* < 2.2 × 10^-16^. Since each BXD line’s singleton mutations should, by definition, be unique to that line, we assumed that each singleton was an independent observation of a particular mutation. However, approximately 50% of each BXD RIL genome is expected to be derived from DBA/2J and 50% is expected to be derived from C57BL/6J; as a result, a pairwise kinship matrix constructed from BXD genotype data will contain non-zero values at essentially every position. To account for kinship between strains in our comparison of singleton fractions, we fit a mixed effects model using the ‘lmekin’ framework from the ‘coxme’ R package. This model predicted C>A singleton fractions as a function of BXD haplotypes at the QTL on chromosome 4, and included the BXD kinship matrix as a random effect term to account for inter-line relatedness. The p-value associated with the ‘haplotype_at_qtl’ fixed effect term remained highly significant (*p* < 2.2 × 10^-16^).

### Comparing mutation spectra in BXDs to TOY-KO triple knockout germline mutation spectra

Exome sequencing was previously performed on a large pedigree of mice with triple knockouts of *Mth1, Mutyh*, and *Ogg1* (in a C57BL/6J background) in order to identify *de novo* germline mutations in mice lacking base excision repair machinery ^11^. The authors deposited all 263 *de novo* germline mutations observed in these mice in the “Supplementary Data File 1” associated with their manuscript. For each mutation, the authors report the reference and alternate alleles, as well as 50bp of flanking sequence up-and downstream of the mutation. We used this information to construct a 3-mer mutation type (ACA>AAA, ACT>AAT, etc.) for each C>A mutation, and then correlated the fractions of each of the 252 C>A 3-mers in the TOY-KO dataset with the enrichments of 3-mer C>A mutation types in BXDs with *D* vs. *B* haplotypes at the QTL on chromosome 4.

### Identifying COSMIC mutation signatures that explain mutation spectrum differences between mice with *B* and *D* haplotypes at the QTL

To uncover more of the genetic etiology of the C>A QTL we observed on chromosome 4, we used a tool called SigProfilerExtractor ^12^ to decompose the mutation spectra of BXD autosomal singletons into distinct sets of COSMIC mutation signatures. In every BXD line, we counted the numbers of singleton mutations belonging to each of the 96 possible 3-mer mutation types (AAA>ATA, AAA>ACA, etc.). We then ran the ‘sigProfilerExtractor’ command on the file containing per-strain counts of each mutation type, specifying ‘maximum_signatures=10’,‘nmf_replicates=100’, and ‘opportunity_genome=“mm10”’.

### Comparing mutation spectra in BXDs to COSMIC mutation signatures

We downloaded mutation signature data for the SBS18 signature from the Catalogue of Somatic Mutations in Cancer (COSMIC) web page using the “Download signature in numerical form” button: https://cancer.sanger.ac.uk/cosmic/signatures/SBS/SBS18.tt. We then correlated the abundances of C>A mutations in all 16 possible 3-mer contexts in the signature with the enrichment of each corresponding 3-mer C>A mutation observed in BXDs with *D* vs. *B* haplotypes at the QTL on chromosome 4.

### Generating phylogenetic comparisons of MUTYH protein sequences

We constructed an alignment of *Mutyh* amino acid sequences for the 9 species shown in **Figure 4** using the web-based Constraint-based Multiple Alignment Tool (COBALT) ^13^and the following NCBI accessions: *Mus musculus* (XP_006503455.1), *Rattus norvegicus* (XP_038965128.1), *Homo sapiens* (XP_011539799.1), *Pan troglodytes* (XP_009454580.1), *Mus caroli* (XP_029332110.1), *Mus pahari* (XP_029395766.1), *Gallus gallus* (XP_004936806.2), *Xenopus tropicalis* (NP_001072831.1), and *Danio rerio* (XP_686698.2). On the COBALT results page, we first downloaded the resulting protein alignment in FASTA format. We then used the “Phylogenetic Tree View” to visualize and download the phylogenetic tree in Newick format.

We then downloaded *Mutyh* coding sequences for each of the above species from the NCBI Nucleotide browser using the following accessions: *Mus musculus* (NM_133250.2), *Rattus norvegicus* (XM_039109200.1), *Homo sapiens* (XM_011541497.3), *Pan troglodytes* (XM_009456305.2), *Mus caroli* (XM_029476250.1), *Mus pahari* (XM_029539906.1), *Gallus gallus* (XM_004936749.3), *Xenopus tropicalis* (NM_001079363.1), and *Danio rerio* (*XM_681606*.*7*). We queried the NCBI Nucleotide database for each accession, used the “Highlight Sequence Features” option to identify the coding sequence, and downloaded the DNA coding sequence in FASTA format.

To visualize both the phylogenetic tree and corresponding *Mutyh* multiple protein sequence alignment, we first reformatted the protein alignment so that it included three separate entries for C57BL/6J, DBA/2J, and the BXD68 line. Specifically, we modified the *Mus musculus* amino acids at positions 5, 24, 69, 312, 313 to create a new DBA/2J sequence, additionally modified the amino acid at position 153 to create a new BXD68 sequence from the DBA/2J sequence, and treated the canonical *Mus musculus* sequence as the C57BL/6J sequence. We reformatted the *Mutyh* coding sequences in the same way, in order to generate three separate entries for *Mus musculus* corresponding to C57BL/6J, DBA/2J, and BXD68.

We then performed a codon-aware multiple sequence alignment of the reformatted coding sequences using the software tool *pal2nal* ^14^. Finally, we visualized the Newick tree and associated multiple protein sequence alignment using the Python API of the *ete3* toolkit ^15^. During this analysis of the protein and coding sequences, we also used the BioPython library ^16^.

### Comparing mutation spectra between groups of Mouse Genomes Project strains

Strain-private substitutions were previously identified in whole genome sequencing data from 29 inbred laboratory mouse strains, which likely represented recent *de novo* germline mutations that occurred in breeding colonies of these strains ^17^. To compare the mutation spectra between various subsets of these strains (grouped by the number of *Mutyh* missense mutations they possessed), we first downloaded “Supplementary Data File 1” from the associated manuscript ^17^. We used data from “Table S3,” which includes both the counts of each mutation type in each strain, as well as the total number of A, T, C, and G base pairs that passed filtering criteria in each strain. Assuming we were comparing the spectra between group “A” and group “B,” for each mutation type we summed the total number of mutations of that type in group “A” and in group “B,” and we collapsed strand complements in our counts (i.e., C>T and G>A are considered to be the same mutation type). Note that the prior report ^17^ does not differentiate summarized counts of C>T mutations into CpG and non-CpG mutations, so our reanalysis of their data does not include the CpG>TpG mutation spectrum category that is part of our BXD analysis. We then summed the total number of callable base pairs corresponding to the reference nucleotide and its complement in group “A” and group “B.” As an example, if we were comparing the counts of C>T mutations between two groups, we summed the counts of callable “C” and “G” nucleotides in each group. We then adjusted the counts of each mutation type in either group “A” or “B” as follows. If the number of callable base pairs was larger in group “B,” we calculated the ratio of callable base pairs between “A” and “B.” We expected that if there were more callable base pairs in a group, then the number of mutations observed in that group might be higher simply by virtue of there being more mutable nucleotides. Therefore, we then multiplied this ratio by the count of mutations in group “B” in order to scale the “B” mutation count down. If the number of callable base pairs were higher in “A,” we performed the same scaling to the counts of mutations in “A.” For each mutation type *i*, we then performed a Chi-square test of independence using a contingency table of four values: 1) the scaled count of mutations of type *i* in group “A”, 2) the scaled count of singleton mutations of type *i* in group “B”, 3) the sum of scaled counts of mutations <u>not</u> of type *i* in group “A”, 4) the sum of scaled counts of mutations <u>not</u> of type *i* in group “B.”

## Main Text Statements

## Acknowledgments

We thank Beth Dumont, Melissa Gymrek, Milad Mortazavi, Andrew Clark, Daphna Rothschild, Uma Arora, Mikhail Maksimov, Hao Chen, and Jonathan Sebat for contributing helpful feedback as part of the BXD Genome Sequencing Consortium. We also thank Molly Przeworski for providing comments on a manuscript draft, Yu-Yu Ren for assistance with preliminary genotype calling, and members of the Harris and Pritchard labs for additional helpful discussions. We thank members of the staff of HudsonAlpha—Dr. Sean Levy and team—for DNA library preparation and sequencing, and for providing us with great support on data transfer. We thank staff at the UT ISAAC facility for storage and processing of all sequence-associated data files. We thank Arthur Centeno for assisting with the upload of phenotype data to GeneNetwork. Finally, we thank Cat Lutz, Alicia Valenzuela at The Jackson Laboratory and Jesse Ingels at UTHSC for DNA sample acquisition, handling, and assistance. We acknowledge support from NIH T32 Postdoctoral Training Grant 5T32HG000035-25 (to TAS), a University of Tennessee Center for Integrative and Translational Genomics grant (to RWW, DGA, LL), NIH Biological Mechanisms of Healthy Aging Training Grant T32AG066574 (to ACB), NIH NIDA Grant P50DA037844 (to AAP), NIH NIGMS Grant 1R01GM123489 (to RWW), NIH NIDA Grant P30 DA044223 (to RWW), NIH R01 HG008140 (to JKP), NIH NIGMS Grant 1R35GM133428-01 (to KH), a Searle Scholarship (to KH), a Sloan Research Fellowship (to KH), and a Pew Biomedical Scholarship (to KH).

## Author contributions

TAS, DGA, AAP, RWW, JKP, and KH contributed to study conceptualization and design. DGA, LL, JKP, AAP and RWW contributed to data curation, including maintenance of the BXD family, DNA sequencing, and data warehousing. TAS, DGA, KH, and ACB contributed to formal data analysis. TAS and KH wrote the original draft of the manuscript. TAS, DGA, AAP, RWW, JKP, and KH contributed to review and editing of the manuscript.

## Additional Information

Correspondence and requests for materials should be addressed to Dr. Kelley Harris.

## Data Availability Statement

BXD mutations and other data files necessary to reproduce the manuscript are available alongside code at https://github.com/tomsasani/bxd_mutator_manuscript (archived at Zenodo, DOI 10.5281/zenodo.5822598). A VCF file containing all variant calls from the sequenced BXDs is being uploaded to the European Variation Archive.

## Code Availability Statement

All code used for data analysis and figure generation is deposited at https://github.com/tomsasani/bxd_mutator_manuscript (archived at Zenodo, DOI 10.5281/zenodo.5822598).

