## Extended Data for "A natural mutator allele shapes mutation spectrum variation in mice"

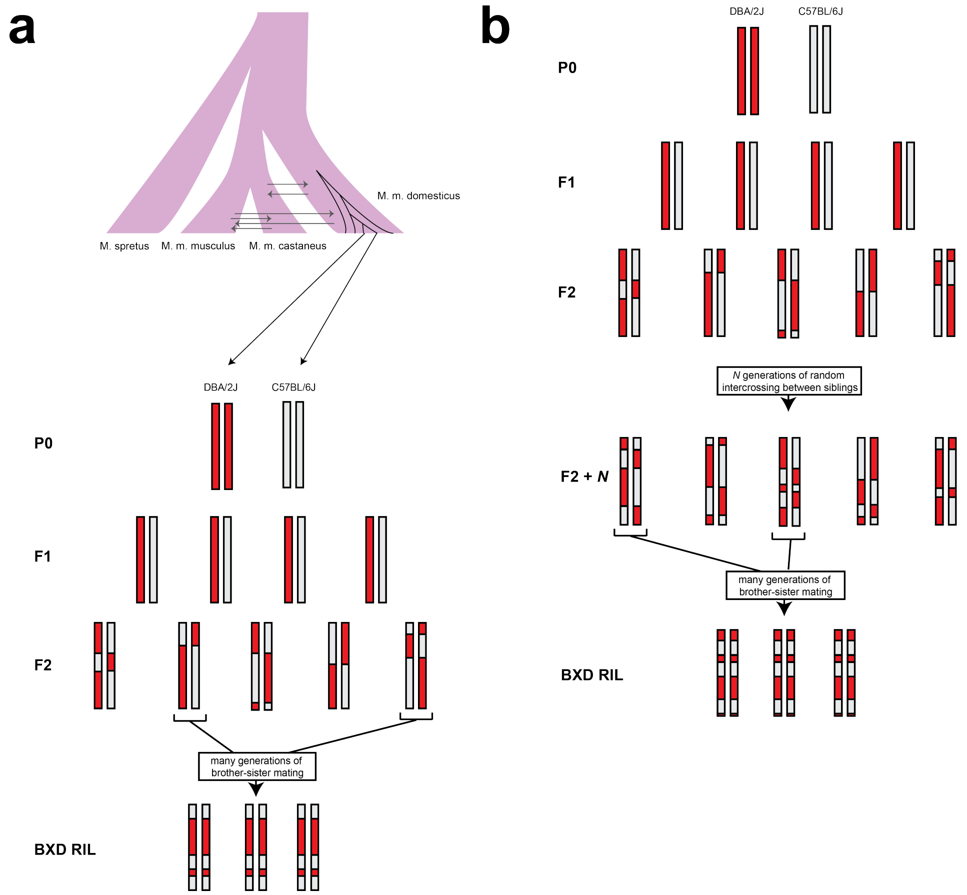


**Extended Data Figure 1: Cross design for BXD RIL construction**

(a) BXDs derived from F2 crosses were subject to many generations of brother-sister mating in order to generate inbred RILs. The genomes of the parents of the BXD crosses (DBA/2J and C57BL/6J) are largely derived from *Mus musculus domesticus*.

(b) To generate advanced intercross lines (AILs), pseudo-random pairs of F2 animals were crossed for *N* generations, and then subject to many generations of brother-sister mating to generate inbred RILs.

a

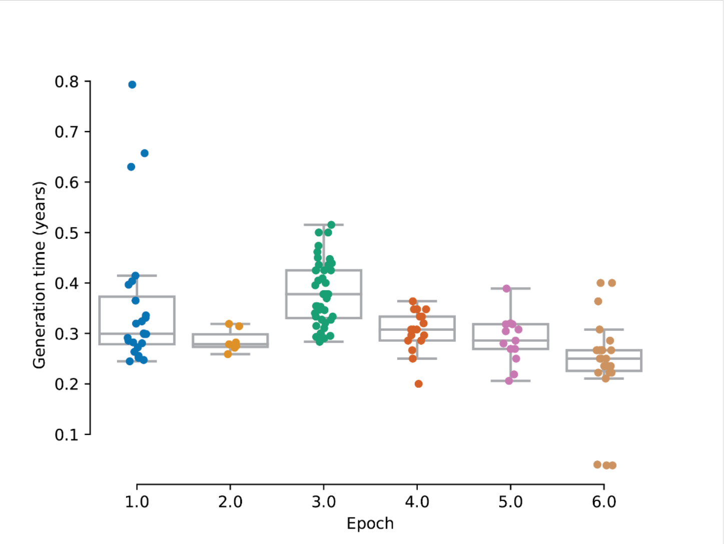


b

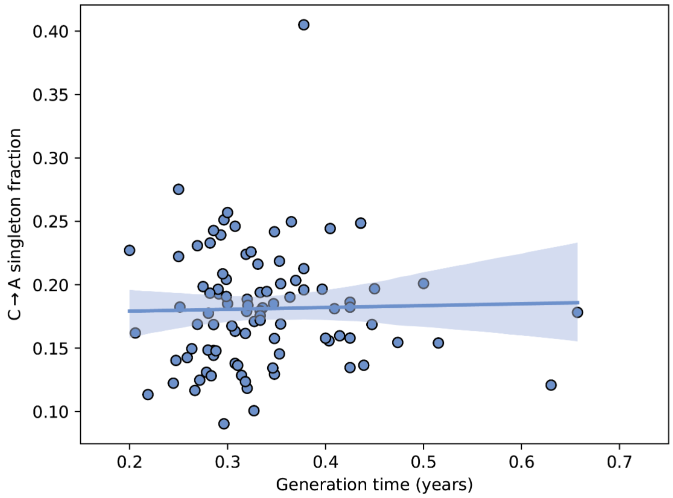


**Extended Data Figure 2: Generation times in the BXD lines**

a) The elapsed time since the founding of each BXD line was calculated by subtracting its initial breeding date from 2017. The elapsed number of years was then divided by the cumulative number of generations of inbreeding undergone by the line, to obtain an estimate of the line's generation time in years.

b) A linear model predicting the C>A singleton fraction in each line as a function of both generation time and the line's epoch of origin was fit using the BXD singleton mutations. C>A fractions were not significantly correlated with generation time (ANOVA $p$ = 0.68).

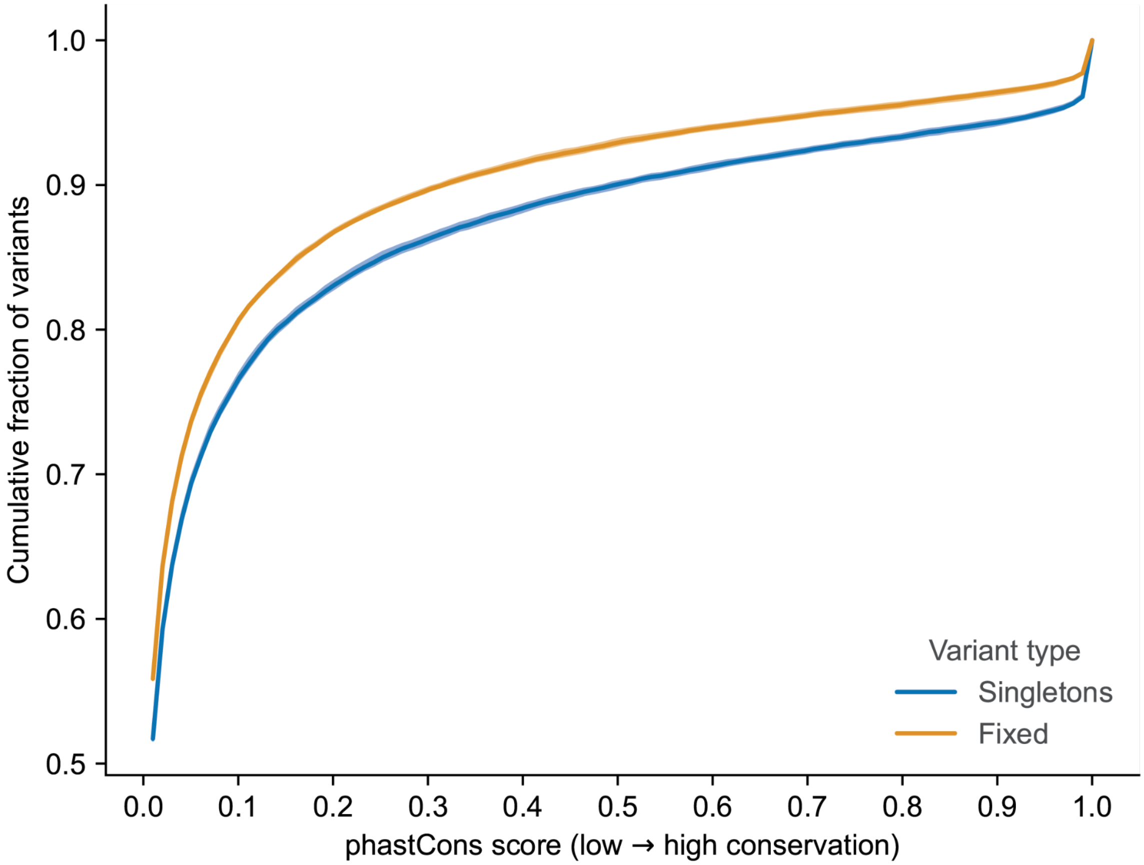


**Extended Data Figure 3: Singletons are enriched in highly conserved regions of the genome**
Cumulative distributions of phastCons conservation probabilities in either singletons (*n* = 47,659) or "fixed" variants (*n* = 81,186). The latter were present in a founder genome and inherited by all BXDs with the founder's haplotype at that site. *P*-value of Kolmogorov-Smirnov test comparing distributions of phastCons scores is 3.8 x $10^{-53}$. Shaded area around each line indicates the bootstrap 95% confidence interval.

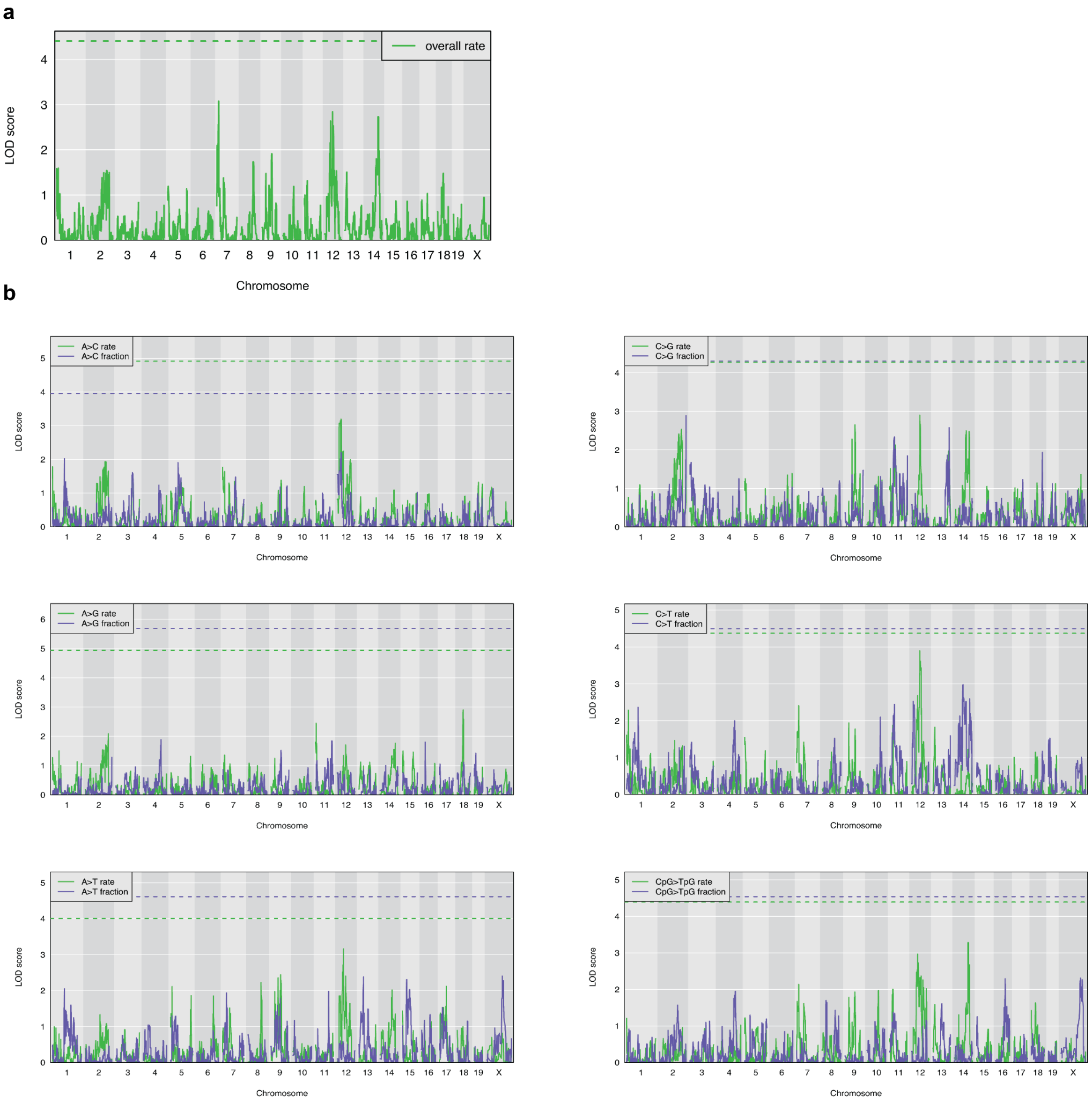


**Extended Data Figure 4: Results of QTL scans for other mutation rate phenotypes**

a) Using the same lines and covariates as described in the Methods, a QTL scan was performed for the overall mutation rate of each line. The green dashed line indicates the genome-wide significance threshold using 1,000 permutations (alpha = 0.05/15).

b) Using the same lines and covariates as described in the Methods, QTL scans were performed for the rates and fractions of all mutation types other than C>A. Green and blue dashed lines indicate the genome-wide significance thresholds for the rate and fraction scans, respectively, using 1,000 permutations (alpha = 0.05/15).

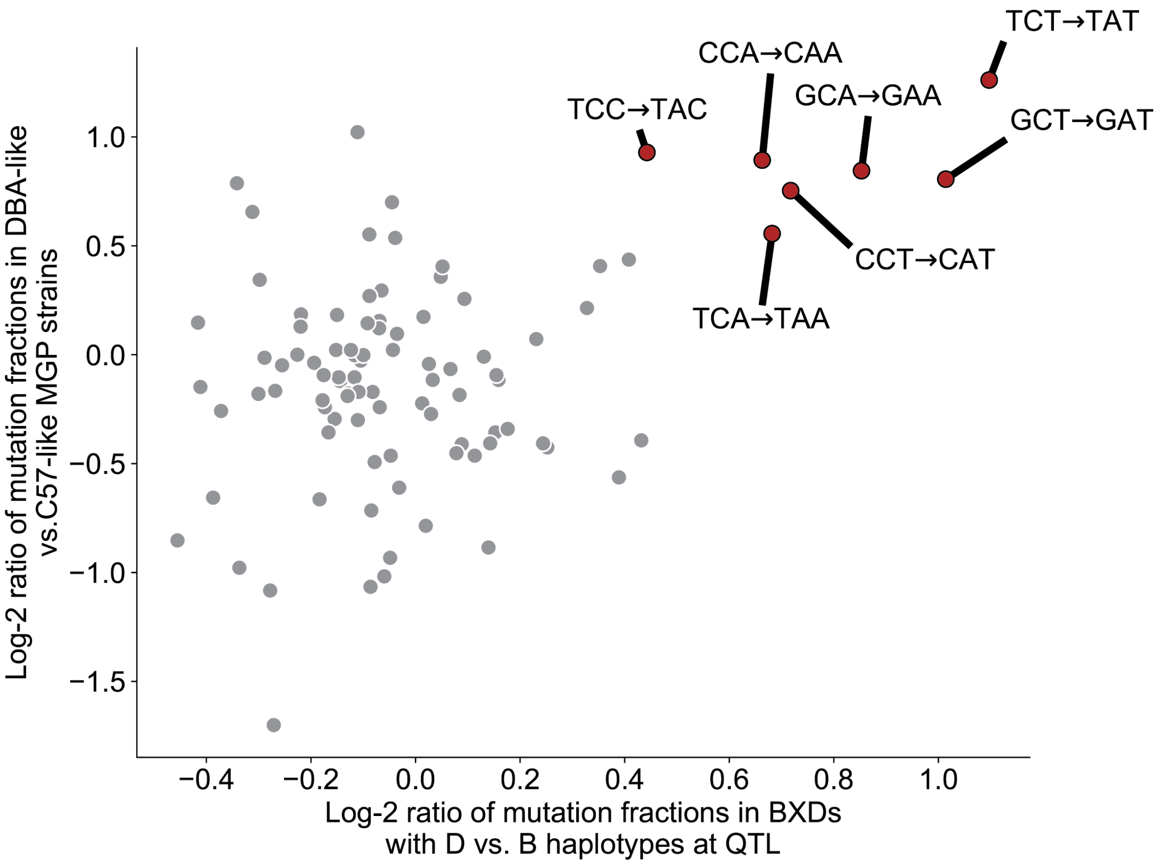


**Extended Data Figure 5: 3-mer mutation sequence contexts enriched in DBA/2J-like Mouse Genomes Project strains**

Log-2 ratios of 3-mer singleton mutation fractions in BXD strains with *D* vs. *B* haplotypes at the QTL on chromosome 4 were correlated with the log-2 ratios of 3-mer strain-private mutation fractions in Sanger Mouse Genomes Project strains that are *D*-like vs. *B*-like. Mutation types significantly enriched in BXDs with *D* haplotypes are colored red, outlined in black and labeled.

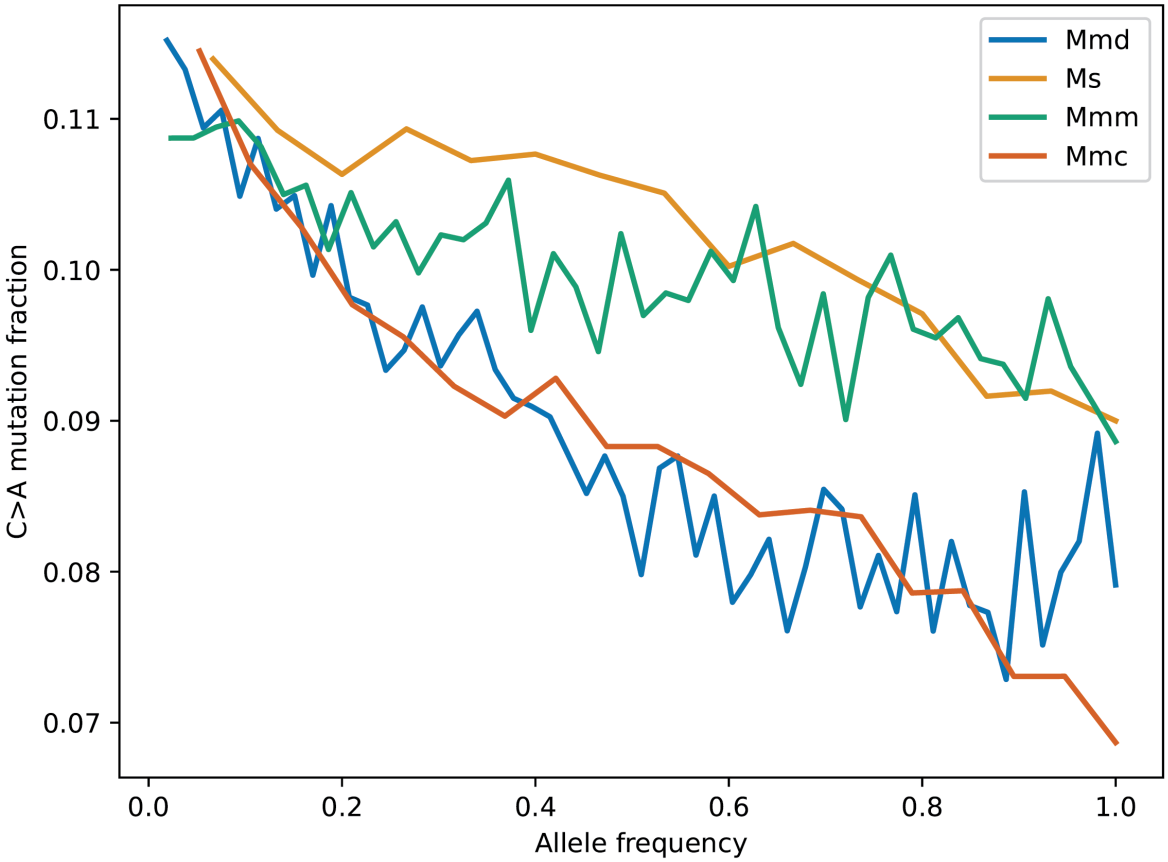


**Extended Data Figure 6: Site frequency spectrum of C>A mutations in the four *Mus* species/subspecies on chromosome 4**

Site frequency spectrum of C>A mutations for four *Mus* species or *Mus musculus* subspecies was generated using a dataset of wild mouse sequencing data from a prior report and a polarized version of the GRCm38/mm10 reference genome. Mmd is *Mus musculus domesticus,* Ms is *Mus spretus,* Mmm is *Mus musculus musculus,* and Mmc is *Mus musculus castaneus.*

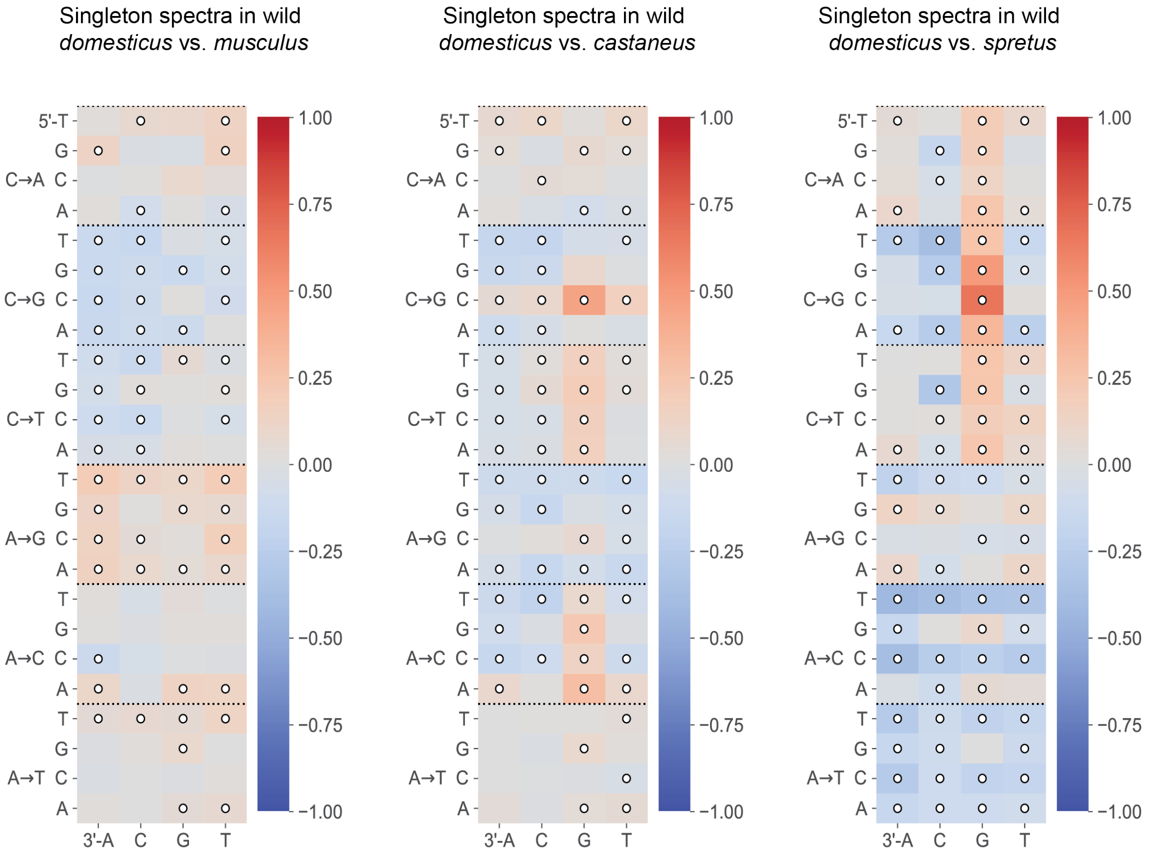


**Extended Data Figure 7: Comparisons of singleton spectra between wild *M.m.domesticus* and other wild species**

Log-2 ratios of singleton fractions of each 3-mer mutation type in *Mus musculus domesticus*, compared to three other wild subspecies or species of *Mus*. Comparisons with Chi-square test of independence *p*-values < 0.05/96 are annotated with white circles.

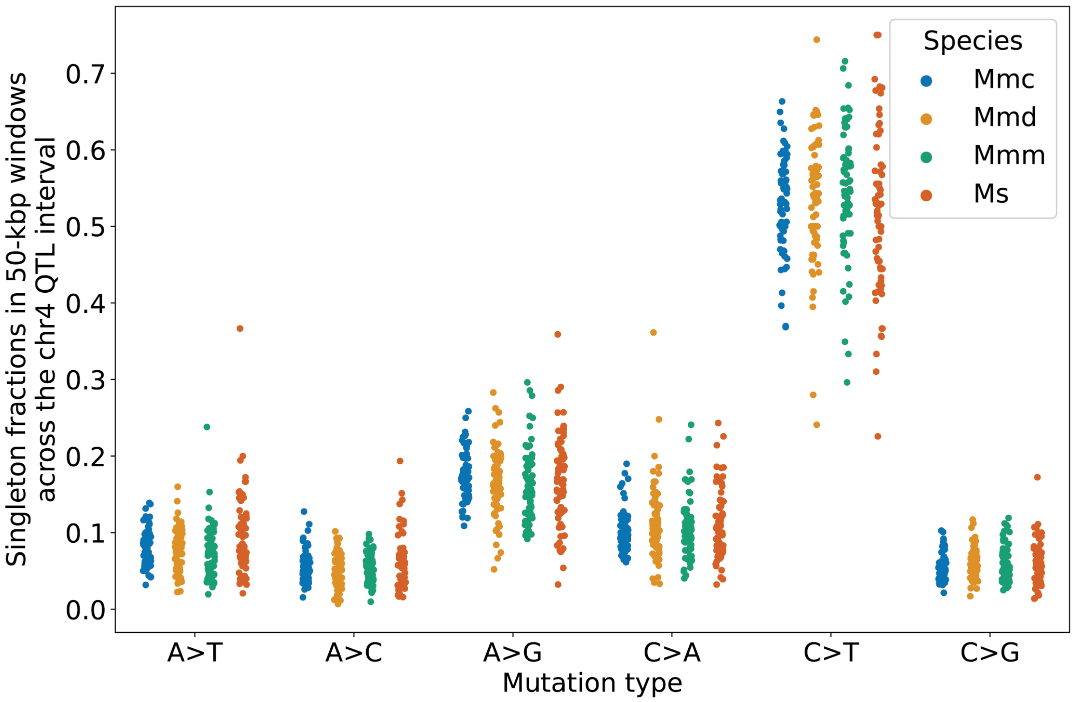


**Extended Data Figure 8: Comparisons of singleton spectra between wild species or subpspecies surrounding *Mutyh***

Singleton fractions of each mutation type in each wild species or subspecies were computed in 50-kilobase pair windows in the QTL interval surrounding *Mutyh* (114.8 Mbp to 118.3 Mbp). The median absolute deviations of C>A fractions in the species or subspecies were: 0.00985 (*Mmc*), 0.0195 (*Mmd*), 0.0155 (*Mmm*), and 0.0214 (*Ms*).

**Extended Data Table 1: BXD RILs used in this analysis**

| Epoch | Year founded | Cross strategy | Strains with available whole-genome sequencing data | Strains in this analysis |
|---|---|---|---|---|
| 1 | 1971 | F2 | 25 | 21 |
| 2 | 1990s (approx.) | F2 | 7 | 7 |
| 3 | late 1990s (approx.) | Advanced intercross | 49 | 38 |
| 4 | 2008 | F2 | 23 | 16 |
| 5 | 2010 | Advanced intercross | 19 | 12 |
| 6 | 2014 | F2 | 30 | 0 |
| Total | - | - | 153 | 94 |
