## Supplementary Information for "A natural mutator allele shapes mutation spectrum variation in mice"


**Criteria for including BXD strains in downstream analyses** **1**

**Identifying haplotypes shared IBD between the BXD RILs and the parental DBA/2J and C57BL/6J strains** **2**

**Identifying fixed variants for comparison to singletons** **3**

**Adjusting singleton counts by the duration of inbreeding in each strain** **4**

**Comparing BXD mutation spectra to previous studies of germline mutation in mice** **5**

**Fine-mapping the QTL to a specific protein-coding gene** **5**

**Identifying correlations between QTL markers and the expression of other genes** **6**

**Identifying fixed structural variant differences between DBA/2J and C57BL/6J within the QTL interval** **8**

**Estimating the strength of the prior association between Mutyh and SBS18 in the scientific literature** **9**

**Generating mutation spectra from wild mouse genomes** **10**

**Scans for natural selection on the mutator locus** **11**

**Generating site frequency spectra of C>A and other mutation types in wild mice** **13**

**Estimating the strength of selection for the B antimutator allele** **14**

**Table 1: GeneNetwork IDs of mutagenesis-related phenotypes that were used for QTL scans** **16**

**Table 2: Numbers of singletons in each BXD line assigned to each COSMIC mutation signature by SigProfilerExtractor** **17**

**References** **20**


### Criteria for including BXD strains in downstream analyses

In this study, our mutation rate estimates were derived from the cumulative number of homozygous autosomal singletons that had accumulated in each strain between its founding and its sequencing. During inbreeding, each generation of brother-sister mating halves the expected heterozygosity of an RIL; a line is predicted to be 99.8% homozygous after 20 generations of inbreeding [1,2]. Thus, if a mutator locus were responsible for influencing the mutation rate or spectrum, that locus would have greater than a 99% chance of being fixed or lost after 20 generations. Moreover, each generation of inbreeding provides an opportunity for heterozygous *de novo* germline mutations to be either fixed or lost. To ensure that any potential mutator loci had been homozygous for a sufficient number of generations to influence the mutation rate or spectrum in each RIL, and that the strains we analyzed had been inbred for long enough to accumulate a substantial number of homozygous *de novo* mutations, we removed BXD RILs that had been inbred for fewer than 20 generations from our analyses. This effectively removed all RILs from epoch 6, the most recent epoch of the BXD.

Also, during construction of the BXDs, 21 RILs were backcrossed to either an inbred DBA/2J or C57BL/6J animal at some point during inbreeding, usually in order to rescue those RILs in cases of severe inbreeding depression [3]. During each generation of backcrossing, a RIL could accumulate new mutations from either the DBA/2J or C57BL/6J parent, as well as from the RIL parent. Additionally, assuming a strain had been inbred prior to backcrossing, each generation of backcrossing is expected to remove half of any existing singleton variants that had accumulated during the prior period of inbreeding in that line. Therefore, to avoid the complexity of accounting for founder-derived singleton mutations and the loss of existing singletons in our estimates of generation times, we removed BXD RILs that had undergone backcrossing from our QTL analyses. Finally, a small number of BXD RILs are nearly or completely isogenic (if two mice from the same line were sequenced, or if the same animal was sequenced twice); these strains were not included in any analyses.

#### Identifying haplotypes shared IBD between the BXD RILs and the parental DBA/2J and C57BL/6J strains

For each chromosome in each recombinant inbred line, we used a Hidden Markov Model (HMM) to identify haplotype tracts that were likely identical-by-descent (IBD) with either of the two founder lines. First, we iterated over every variant in the joint-genotyped BXD VCF file; we only considered sites where one of the two founder strains (either DBA/2J or C57BL/6J) was homozygous for the alternate allele, and the other strain was homozygous for the reference allele. We further required the genotype of each founder to be supported by at least 10 sequencing reads, and the Phred-scaled genotype quality of each founder genotype to be at least 20. We assumed that these sites represented fixed differences between DBA/2J and C57BL/6J. By comparing the genotypes of all BXD RILs to the two founder genotypes at each of these informative fixed differences, we then identified tracts of sites in which RIL genotypes consistently matched one of the two founders. For each chromosome in each BXD RIL, we constructed an array of length $S$, where $S$ was the number of fixed differences between C57BL/6J and DBA/2J across that chromosome. Each element $i$ in this array took one of three values: 0 (if the genotype of the RIL at site $S_i$ did not match the DBA/2J genotype), 1 (if the genotype of the RIL at site $S_i$ matched the DBA/2J genotype), or -1 (if the genotype of the RIL at site $S_i$ was unknown, heterozygous, supported by fewer than 10 reads, or had a Phred-scaled genotype quality < 20). We then built an HMM using *pomegranate* [4] with two states (DBA/2J or C57BL/6J, state and transition probabilities defined in associated code), and inferred long sequences of identical states. These sequences likely represent regions of the genome inherited from one of the two founders.

#### Identifying fixed variants for comparison to singletons

We aimed to compare the distribution of phastCons conservation probabilities in the BXD singletons to a "control" set of variants. We defined these control variants as mutations present in one of the two founder genomes (which passed the same filtering criteria required of singletons), and present in all BXDs that inherited the founder haplotype with the mutation at that site. We expected that these "fixed" variants had occurred well before the construction of the BXDs in DBA/2J or C57BL/6J founder stocks. To ensure that we sampled control variants from the full length of the reference genome sequence, we first used *bedtools* [5] to generate 50-kbp windows across the mm10/GRCm38 reference genome using the "makewindows" subcommand, and then identified all of the fixed mutations in each window. When comparing the distributions of phastCons scores between fixed and singleton mutations, we sampled five fixed mutations from each 50-kbp window in order to obtain a number of fixed mutations that more closely matched the number of singleton mutations.

### Adjusting singleton counts by the duration of inbreeding in each strain

We downloaded a file containing BXD metadata (strain names, provenance, etc.) from a prior manuscript introducing the updated BXD family [3] (available from the cited manuscript as SuppTable1.xlsx). A simplified version of this file, containing only the metadata necessary to reproduce the analyses in this manuscript, is provided in the GitHub repository associated with this manuscript. In this file, each BXD strain is annotated with its "Generation at sequencing," using the Jackson Laboratories Generation Definitions syntax. For example, a strain that was inbred for 25 generations at its original location and subsequently inbred for 20 generations after arriving at the Jackson Laboratories would be annotated as "F25+F20." To assign an "age" in generations to each BXD RIL, we simply summed the number of total generations each strain had been inbred; in the previous example, the strain's age would be 45 total generations.

To estimate each strain's mutation rate per genome per generation, we used a previously described approach that assumes perfect full-sibling mating during inbreeding [6]. Briefly, we divided the count of homozygous singletons in each mouse by the number of generations in which *de novo* germline mutations could have occurred in the strain, and by the number of haploid base pairs that were callable (i.e., were covered by at least 10 sequencing reads in the strain, and not overlapping segmental duplications or simple repeats). To calculate the callable number of base pairs, we used *mosdepth* [7] to count the number of base pairs with at least 10 aligned sequencing reads (of at least quality score 20) in each BXD RIL's BAM file with the following command:

```
mosdepth -n -x -b 1000000 -T 10 -Q 20 $PREFIX $BAM
```

We then used the files produced by this command to count the total number of autosomal base pairs covered by at least 10 high-quality sequencing reads that did not overlap segmental duplications or simple repeat tracks from the UCSC Genome Browser. Threshold files produced by *mosdepth* for each BXD are included at the GitHub repository associated with this manuscript.

### Comparing BXD mutation spectra to previous studies of germline mutation in mice

A recent study estimated germline mutation rates in mice[8] that were generated via a cross between two inbred laboratory strains: C57BL/6J and 129S5. Using these previously published germline mutation data, we estimated the 95% confidence interval bounding the *de novo* C>A mutation fraction to be between 0.119 and 0.167. This 95% CI includes the C>A fraction of the strain-private mutations measured in C57BL/6NJ by Dumont (2019) (0.134), and the average C>A singleton fraction in BXD mice with *B* haplotypes at the QTL (0.141). Additionally, we note that the C>A fraction of strain-private mutations measured in DBA/2J by Dumont (2019) (0.2081) is almost exactly the same as the average C>A singleton fraction in BXD mice with *D* haplotypes at the QTL (0.2084).

### Fine-mapping the QTL to a specific protein-coding gene

To narrow our search for sequence variation underlying the C>A QTL on chromosome 4, we first used the Mouse Genome Informatics (MGI) query tool to identify any protein-coding genes within the QTL. Using the "Genes and Markers Query Form" (http://www.informatics.jax.org/marker), we searched for protein-coding genes within the Bayes

95% credible QTL interval on chromosome 4 (from 114.8 - 118.3 Mbp). A total of 76 protein-coding genes overlapped the QTL. We then used the MGI "Batch Query" tool to determine which of these genes was associated with the following Gene Ontology terms: "DNA replication, "cellular response to DNA damage."

We then asked if any of the protein-coding genes within the QTL interval contained sequence differences between DBA/2J and C57BL/6J. We iterated over every variant within the QTL interval that was homozygous for the alternate allele (or heterozygous, with >= 0.9 allele balance) in either DBA/2J or C57BL/6J and homozygous for the reference allele in the other strain; we then examined the SnpEff annotations associated with each of these variants. Importantly, SnpEff may add more than one annotation to a variant if that variant affects more than one transcript sequence. For each high-quality fixed difference between DBA/2J and C57BL/6J, we asked if the variant was annotated as having MODERATE or HIGH impact on any of the transcript sequences reported in the "ANN" entry added to the INFO field of the variant. We considered any variant with at least one MODERATE or HIGH-impact annotation to be of interest.

#### Identifying correlations between QTL markers and the expression of other genes

We used the GeneNetwork online resource [9] to ask if the QTL for the C>A mutation rate on chromosome 4 could be a result of the QTL harboring expression quantitative trait loci (eQTLs) for the expression of genes involved in DNA repair or genome integrity. Specifically, we searched for associations between the top SNP marker (i.e., the marker with highest LOD score) from the C>A QTL interval on chromosome 4 (rs52263933) and the expression of protein-coding genes within the QTL interval. On the GeneNetwork website (genenetwork.org), we searched the "BXD Genotypes" dataset for rs52263933, after selecting the "DNA Markers and SNPs" type and the "BXD family" group. Using the Trait Data and Analysis page for rs52263933, we then calculated correlations between the marker and expression datasets from a variety of tissues. We limited the results to include the top 100 genes for which expression was correlated with rs52263933, and used "Sample r" as the correlation method. For each tissue, protein-coding genes that had expression values that were significantly ($p < 0.05$) correlated with BXD genotypes at rs52263933 are listed below; those that were annotated with any one of the following Gene Ontology terms are indicated: “cellular response to DNA damage,” “DNA repair,” or “cellular response to oxidative stress.”

**Amygdala**: INIA Amygdala Cohort Affy MoGene 1.0 ST (Mar11) RMA

- Not annotated with relevant GO terms: *Atpaf1, Ipp, Ccdc17, Urod, Eif2b3, Tmem53, Eri3, B4galt2, St3gal3*

**Hematopoietic Stem Cells**: UMCG Stem Cells ILM6v1.1 (Apr09) original

- Not annotated with relevant GO terms: *Faah, Hpdl, Kif2c*
- Annotated with relevant GO terms: *Mutyh*

**Kidney**: Mouse kidney M430v2 Sex Balanced (Aug06) RMA

- Not annotated with relevant GO terms: *Pdzk1ip1, Atpaf1, Faah, Tspan1, Ipp, Rps8, Atp6v0b, Ptprf*

**Liver**: UTHSC BXD Liver RNA-Seq Avg (Oct19) TPM Log2

- Not annotated with relevant GO terms: *Cyp4a32, Atpaf1, Mob3c, Faah, Ccdc17, Urod, Tmem53, Eri3, St3gal3*
- Annotated with relevant GO terms: *Mutyh*

**Gastrointestinal**: UTHSC Mouse BXD Gastrointestinal Affy MoGene 1.0 ST Gene Level (Apr14) RMA

- Not annotated with relevant GO terms: *Stil, Atpaf1, Mobkl2c, Faah, Ccdc17, Zswim5, Tmem53, Atp6v0b, Ptprf*

**Spleen**: UTHSC Affy MoGene 1.0 ST Spleen (Dec10) RMA Exon Level

- Not annotated with relevant GO terms: *Atpaf1, Mobkl2c, Faah, Ipp, Gpbp1l1, Ccdc17, Btbd19, Eri3, Atp6v0b, St3gal3*
- Annotated with relevant GO terms: *Mutyh, Plk3*

#### Identifying fixed structural variant differences between DBA/2J and C57BL/6J within the QTL interval

To determine the potential impact of fixed structural variant (SV) differences between DBA/2J and C57BL/6J on the C>A mutator phenotype we observed in the BXD, we first used the Sanger Mouse Genomes Project variant query tool (https://www.sanger.ac.uk/sanger/Mouse_SnpViewer/rel-1505) to find all structural variants within the QTL interval on chromosome 4 (between 114.8 and 118.3 Mbp). We limited our search to structural variants that were fixed differences between C57BL/6NJ and DBA/2J. A total of 51 SVs met this criterion; we downloaded the structural variants in BED format from the MGP website.

We then downloaded the complete set of mm10 GENCODE VM23 Genes and Gene Predictions from the UCSC Table Browser in GTF format. Using bedtools [5], we intersected the GENCODE VM23 gene file with the BED file containing the structural variants using default parameters, and counted the number of SVs that overlapped the exonic sequences of any genes. Finally, we used the Mouse Genome Informatics Batch Query resource (http://www.informatics.jax.org/batch) to identify the Gene Ontology terms associated with each of the genes overlapping SVs.

Fixed structural differences between *D* and *B* haplotypes overlapped the exonic sequences of two protein-coding genes within the QTL interval (*Cyp4a32* and *Ptch2*), as well as the sequences of two predicted protein-coding genes (*Gm22398* and *Gm12840*). As none of these genes have a function that is directly related to DNA repair, replication, or the maintenance of genome integrity, it is unlikely that these SVs underlie the C>A QTL on chromosome 4.

### Estimating the strength of the prior association between *Mutyh* and SBS18 in the scientific literature

Since the QTL on chromosome 4 contains many differences between the *B* and *D* haplotypes that occur outside the *Mutyh* gene, our argument that *Mutyh* variation is likely to be causal hinges upon a Bayesian line of reasoning. Bayesian statisticians acknowledge that experimental results are rarely interpreted in a vacuum; instead, experimentalists almost always have a prior belief (informed by previous scientific work) that certain results are more likely than others. In our case, we assume that genes associated with DNA replication and repair are *a priori* more likely to harbor mutator alleles than other genes. However, the strength of this prior is mitigated by the knowledge that variation in other types of genes might impact the mutation spectrum, for example, by affecting cell metabolism in a way that increases or decreases the production of mutagenic metabolites.

Our Bayesian prior that *Mutyh* missense mutations underlie the QTL observed on chromosome 4 is augmented by numerous previous studies that have established a clear link between *Mutyh* deficiency (in particular, missense mutations in *Mutyh*) and C>A mutagenesis. An early report [10] surveyed colorectal tumors from siblings affected with colorectal cancer, and found 18 somatic inactivating mutations in the APC gene. The authors noted that 15 of these 18 inactivating mutations were C>A transversions, and further discovered that all of the affected siblings were compound heterozygous for missense mutations in *MUTYH*. Another early report [11] found that in 16 individuals with either adenomas or polyposis, in addition to biallelic missense mutations in *MUTYH*, all somatic APC mutations were C>A transversions. More recently, mutation signature analyses have uncovered a clear link between germline *MUTYH* mutations and specific C>A dominated mutation signatures, such as SBS18 and SBS36 [12]. For example, one study of 498 colorectal tumors found that attribution of at least 30% of the somatic mutation load to SBS18/SBS36 was 100% predictive of the presence of inherited pathogenic biallelic *Mutyh* missense variants [13]. Another study found that the SBS18 mutation signature was exclusively present in colorectal tumors from patients with pathogenic missense or nonsense mutations in *MUTYH*, and comprised up to 70% of all somatic mutations in these samples [14].

To our knowledge, the only other genes implicated by the literature in SBS18 mutagenesis are *OGG1* (a direct interaction partner of *MUTYH* [15]), the transcription factor *RUNX1* (which regulates *OGG1* [16,17]), and the RNA editing enzyme *APOBEC* [18].

To bibliometrically quantify the prior association between SBS18 and *Mutyh* in the biomedical literature, we also queried Google Scholar for all instances of the "SBS18" term. A total of 192 publications reference SBS18, and 47 of them (nearly 24%) also reference *Mutyh*. Coupled with the biochemical knowledge that MUTYH plays a direct role in repairing 8-oxoguanine lesions and preventing C>A mutations during DNA replication [19], this amounts to a clear prior expectation that *Mutyh* missense mutations likely underlie the C>A QTL on chromosome 4 in the BXD.

### Generating mutation spectra from wild mouse genomes

To identify singleton variants within each wild *Mus* species or subspecies, we analyzed a VCF file containing variant calls for 67 wild mice from four *Mus* species/subspecies from [20]. We iterated over all autosomal variants in the VCF and limited our search to single-nucleotide variants. For each variant, we examined each species or subspecies (*Mus musculus domesticus, Mus musculus castaneus, Mus musculus musculus,* or Mus *spretus*) separately, and considered singleton variants to be mutations present in only one sequenced sample from a particular subspecies. As many of the wild mice are naturally inbred, we allowed singletons to be heterozygous or homozygous in a sample; we limited to sites where the singleton mutation represented an alternate allele, and all other samples were homozygous for reference alleles at the site. However, because we examined each subspecies separately, we included singletons that were observed across subspecies. For example, if one of the 27 *domesticus* samples had a variant at a particular site, and one of the 22 *musculus* samples had the same variant, we considered the variant to be a singleton in both subspecies (and assumed that the singleton occurred independently in both subspecies). We required singletons to be supported by at least 10 sequencing reads, to have a Phred-scaled genotype quality of at least 20, and for heterozygous singletons to have an allele balance (fraction of reads supporting the ALT allele) between 0.25 and 0.75. We excluded singletons that were observed within either segmental duplications or simple repeats (downloaded from the UCSC Table Browser in mm10 coordinates), and singletons that occurred at nucleotides with a phastCons probability of conservation > 0.05.

### Scans for natural selection on the mutator locus

We utilized a number of software tools and statistical tests to detect signals of natural selection on the locus identified by our QTL analysis on chromosome 4. Using the *ete3* toolkit [21], we performed two tests for positive selection on the *Mutyh* gene across a clade of 10 rodent species: *Mus musculus, Rattus norvegicus*, *Mus caroli*, *Mus pahari*, *Meriones unguiculatus*, *Mesocricetus auratus*, *Peromyscus maniculatus bairdii*, *Onychomys torridus*, *Microtus oregoni*, and *Microtus ochrogaster*. In both analyses, we first generated an alignment of MUTYH protein sequences across the 10 rodent species using the web-based COBALT [22] tool, using the following accessions: *Mus musculus* (XP_006503455.1), *Rattus norvegicus* (XP_038965128.1), *Mus caroli* (XP_029332110.1), *Mus pahari* (XP_029395766.1), *Mesocricetus auratus* (XP_012970297.1), *Peromyscus maniculatus bairdii* (XP_006986619.1), *Onychomys torridus* (XP_036033228.1), *Meriones unguiculatus* (XP_021489104.1), *Microtus oregoni* (XP_041509038.1), and *Microtus ochrogaster* (XP_005370081.1). We downloaded both the aligned protein sequences in FASTA format and the COBALT phylogenetic tree in Newick format. We additionally downloaded the *Mutyh* DNA coding sequences for each of these species from the NCBI Nucleotide browser, and generated a codon-aware alignment of the coding sequences using *pal2nal* [23]. Using the *ete3 evol* toolkit's wrapper around PAML [24], we then performed likelihood ratio tests (LRT) between two pairs of models in order to test for positive selection on sites in *Mutyh*: M2 vs. M1 and M8 vs M7. Neither LRT indicated that the null model should be rejected ($p$ = 1.0 and $p$ = 0.87, respectively).

We also uploaded the codon-aware alignment to the Datamonkey server [25,26] and used the BUSTED tool [27] to detect whether there was evidence of gene-wide episodic diversifying selection on at least one site on at least one branch of the rodent clade. This analysis did not return significant evidence for selection on *Mutyh* in the clade (BUSTED LRT $p$ = 0.5).

To detect signatures of selective sweeps using site frequency spectra, we also analyzed previously published wild mouse genomes [20] using SweeD (v3.2.1) [28]. We first generated site frequency spectra separately for the *Mus musculus musculus, Mus musculus domesticus,* and *Mus musculus castaneus* populations on chromosome 4. Next, we ran the following SweeD command for each subpopulation, using its own site frequency distribution:

```
./SweeD -name Mmd -input /path/to/Mmd/sfs/data -grid 1000
./SweeD -name Mmc -input /path/to/Mmc/sfs/data -grid 1000
./SweeD -name Mmm -input /path/to/Mmm/sfs/data -grid 1000
```

SweeD did not return a significant likelihood ratio within the QTL interval on chromosome 4 (114.8 Mbp to 118.3 Mbp) that indicated evidence for positive selection on a haplotype in any of the *Mus* subspecies.

Finally, using the wild mouse data [20] we performed a McDonald-Kreitman test [29] to determine whether there was a significant difference in the numbers of fixed and polymorphic substitutions that were either synonymous or nonsynonymous in *Mutyh*. We used SnpEff [30] to annotate each of the wild variants in *Mutyh* with its predicted impact on the amino acid sequence of MUTYH using the following command:

```
java -Xmx16g -jar /path/to/snpeff/jarfile GRCm38.86 /path/to/wild/vcf
> /path/to/uncompressed/output/vcf
```

and subsequently determined whether each variant was fixed or polymorphic in a particular *Mus* subspecies. We then generated a contingency table using the counts of fixed and polymorphic mutations predicted to be synonymous or non-synonymous by SnpEff; we required that these mutations were covered by at least 10 sequencing reads and had a Phred-scaled genotype quality of at least 20, and removed sites that overlapped annotated segmental duplications or simple repeats in the mm10/GRCm38 genome. We considered any mutation that was homozygous for an alternate allele in all members of a particular subspecies, and homozygous for the reference allele in all other wild samples, to be fixed; if a mutation was not homozygous for an alternate allele in all members of a particular subspecies, or if it was segregating in any other subspecies, we considered it to be polymorphic. We then performed a Chi-squared test of independence to determine if the ratio of non-synonymous to synonymous mutations was significantly different for fixed or polymorphic substitutions (Chi-square $p$ = 1.0).

### Generating site frequency spectra of C>A and other mutation types in wild mice

To generate a site frequency spectrum for C>A mutations in the wild mice from Harr et al. [20], we first used *bcftools* [31] v1.12 to remove sites from the wild mouse VCF where all samples were fixed for the same allele, as well as sites that overlapped regions of the mm10/GRCm38 reference genome annotated as being simple repeats or segmental duplications. We then used *est-sfs* [32] v2.03 to predict the most likely ancestral allele at each variant site in the wild mouse VCF. We treated *Mus mus domesticus* as the focal population, used *Mus musculus musculus* and *Mus spretus* as outgroups one and two, respectively, and parametrized *est-sfs* using the Kimura 2-parameter model (`model 1`). We considered sites at which the probability of the major allele being ancestral was between 0.1 and 0.9 to be "ambiguous," and used *bedtools* *maskfasta* [5] to "mask" those sites in the reference genome by modifying them to be "N"s. However, if the probability of the major allele being ancestral at a particular site was >= 0.9, we modified the reference genome sequence at that site to be the major allele; and if the probability of the major allele being ancestral at the site was <= 0.1, we modified the reference genome sequence at that site to be the minor allele instead. We then used the *mutyper variants* [33] subcommand (*mutyper* v0.5.0) to annotate the wild mouse VCF INFO field with the modified, "ancestral" reference sequence, and the *mutyper ksfs* command to generate a k-SFS (i.e., a site frequency spectrum for each 1-mer mutation type) separately for each subpopulation of wild mice (*Mus mus musculus, Mus mus domesticus, Mus mus castaneus,* and *Mus spretus*). The k-SFS in each wild subpopulation was therefore calculated using only confidently polarized sites.

### Estimating the strength of selection for the *B* antimutator allele

Population genetic theory [34] suggests that the selective disadvantage of a mutator allele is equal to $2s\Delta U$ (assuming the mutator is not completely recessive), where *s* is the mean selection coefficient for a deleterious mutations and $\Delta U$ is the increased deleterious mutation load caused by the mutator allele. The product of $\Delta U$ and *s* is then multiplied by 2 to reflect the fact that each new deleterious mutation is linked to the mutator for an average of two generations. Because BXD lines homozygous for the *B* haplotype avoid about 1.5 mutations per haploid genome per generation compared to lines homozygous for the *D* haplotype, coding sequence comprises approximately 2% of the mouse genome, and assuming that nonsynonymous coding mutations have an average selection coefficient of $-5 \times 10^{-3}$ [35], we estimated that the *B* allele should enjoy a fitness advantage of about $2 * (1.5 * 0.02) * 5 \times 10^{-3} = 3 \times 10^{-4}$ over the *D* allele if additive, and $6 \times 10^{-4}$ if dominant. Assuming a wild mouse effective population size of approximately $N = 5 \times 10^4$, this would make $2Ns = 30$ or 60, comfortably exceeding the threshold of $2Ns = 1$ at which a mutator is expected to segregate neutrally[36]. As the BXD are purposefully inbred and maintained in a highly controlled laboratory environment, and because we may rely on somewhat simplistic assumptions and parameter estimates, our estimates of selection may not translate to wild species and subspecies of *Mus*; however, our results may provide useful parameters for estimating the effects of mutator alleles in natural populations. Other factors might also reduce the selective advantage of the *B* antimutator alleles; these include the fact that the allele must reach mutation-selection balance with respect to its reduced mutation load (ΔU) before reaching its full selective advantage [37,38], and any potential cost of replication associated with the mutator allele [38].

Table 1: GeneNetwork IDs of mutagenesis-related phenotypes that were used for QTL scans

| **Phenotype** | **GeneNetwork ID** | **# of mutations in estimate** |
|---|---|---|
| C>A singleton fraction | BXD_24430 | 11,728 |
| C>T singleton fraction | BXD_24431 | 19,868 |
| C>G singleton fraction | BXD_24432 | 4,045 |
| A>C singleton fraction | BXD_24433 | 4,022 |
| A>T singleton fraction | BXD_24434 | 5,731 |
| A>G singleton fraction | BXD_24435 | 9,005 |
| CpG>TpG singleton fraction | BXD_24436 | 9,515 |
| C>A mutation rate (per base pair, per generation) | BXD_24437 | 11,728 |
| C>T mutation rate (per base pair, per generation) | BXD_24438 | 19,868 |
| C>G mutation rate (per base pair, per generation) | BXD_24439 | 4,045 |
| A>C mutation rate (per base pair, per generation) | BXD_24440 | 4,022 |
| A>T mutation rate (per base pair, per generation) | BXD_24441 | 5,731 |
| A>G mutation rate (per base pair, per generation) | BXD_24442 | 9,005 |
| CpG>TpG mutation rate (per base pair, per generation) | BXD_24443 | 9,515 |
| Overall mutation rate (per base pair, per generation) | BXD_24444 | 63,914 |

Table 2: Numbers of singletons in each BXD line assigned to each COSMIC mutation signature by SigProfilerExtractor

| Samples | SBS1 | SBS5 | SBS18 | SBS30 |
|---|---|---|---|---|
| BXD001_TyJ_0361 | 193 | 674 | 304 | 220 |
| BXD002_RwwJ_0430 | 291 | 1210 | 0 | 317 |
| BXD006_TyJ_0474 | 276 | 1095 | 0 | 305 |
| BXD008_TyJ_0372 | 99 | 567 | 0 | 228 |
| BXD009_TyJ_0383 | 187 | 780 | 283 | 203 |
| BXD011_TyJ_0368 | 249 | 1214 | 0 | 380 |
| BXD013_TyJ_0410 | 175 | 735 | 0 | 236 |
| BXD016_TyJ_0393 | 290 | 1101 | 466 | 402 |
| BXD024_TyJ_0347 | 170 | 731 | 0 | 197 |
| BXD028_TyJ_0346 | 136 | 621 | 224 | 205 |
| BXD031_TyJ_0364 | 293 | 987 | 0 | 339 |
| BXD032_TyJ_0415 | 202 | 701 | 280 | 271 |
| BXD034_TyJ_0356 | 126 | 418 | 0 | 151 |
| BXD036_TyJ_0369 | 56 | 258 | 119 | 106 |
| BXD038_TyJ_0453 | 111 | 468 | 0 | 106 |
| BXD045_RwwJ_0486 | 90 | 304 | 128 | 110 |
| BXD049_RwwJ_0397 | 102 | 415 | 0 | 124 |
| BXD050_RwwJ_0419 | 94 | 382 | 128 | 136 |
| BXD051_RwwJ_0408 | 102 | 368 | 152 | 102 |
| BXD056_RwwJ_0421 | 84 | 315 | 154 | 115 |
| BXD064_RwwJ_0483 | 68 | 302 | 90 | 95 |
| BXD073_RwwJ_0438 | 56 | 218 | 64 | 95 |
| BXD086_RwwJ_0403 | 96 | 296 | 184 | 97 |
| BXD090_RwwJ_0395 | 78 | 266 | 182 | 114 |
| BXD095_RwwJ_0445 | 69 | 213 | 63 | 90 |
| BXD098_RwwJ_0345 | 104 | 316 | 106 | 101 |
| BXD100_RwwJ_0469 | 77 | 204 | 126 | 82 |
| BXD101_RwwJ_0475 | 60 | 236 | 79 | 60 |
| BXD102_RwwJ_0406 | 51 | 225 | 70 | 82 |
| BXD111_0424 | 17 | 0 | 39 | 40 |
| BXD113_RwwJ_0365 | 49 | 196 | 122 | 0 |
| BXD122_TyJ_0467 | 36 | 99 | 0 | 30 |
| BXD123_0431 | 36 | 144 | 0 | 43 |

| | | | | |
|---|---|---|---|---|
| BXD124_RwwJ_0454 | 30 | 0 | 34 | 33 |
| BXD125_RwwJ_0472 | 36 | 0 | 83 | 109 |
| BXD128_0492 | 24 | 121 | 0 | 0 |
| BXD12_TyJ_0464 | 241 | 992 | 321 | 267 |
| BXD141_0450 | 34 | 150 | 0 | 0 |
| BXD144_0352 | 35 | 125 | 0 | 41 |
| BXD147_0494 | 25 | 145 | 0 | 0 |
| BXD14_TyJ_0458 | 212 | 787 | 370 | 351 |
| BXD150_0373 | 34 | 86 | 60 | 39 |
| BXD151_0457 | 30 | 114 | 0 | 0 |
| BXD154_RwwJ_0374 | 35 | 115 | 0 | 36 |
| BXD155_0362 | 17 | 80 | 37 | 29 |
| BXD156_0357 | 21 | 91 | 54 | 40 |
| BXD157_0391 | 35 | 97 | 50 | 0 |
| BXD15_TyJ_0392 | 161 | 581 | 422 | 226 |
| BXD160_0427 | 52 | 167 | 0 | 48 |
| BXD161_RwwJ_0456 | 29 | 116 | 67 | 44 |
| BXD168_0466 | 43 | 201 | 0 | 0 |
| BXD169_0426 | 28 | 232 | 0 | 0 |
| BXD170_0437 | 54 | 209 | 0 | 52 |
| BXD171_0484 | 29 | 81 | 34 | 0 |
| BXD172_0380 | 47 | 203 | 0 | 50 |
| BXD177_0490 | 43 | 157 | 0 | 59 |
| BXD178_0342 | 30 | 185 | 0 | 0 |
| BXD180_0353 | 36 | 146 | 0 | 61 |
| BXD184_0452 | 36 | 193 | 0 | 0 |
| BXD186_0416 | 42 | 161 | 0 | 53 |
| BXD18_TyJ_0477 | 232 | 834 | 268 | 289 |
| BXD19_TyJ_0481 | 236 | 1005 | 510 | 327 |
| BXD21_TyJ_0432 | 116 | 506 | 174 | 147 |
| BXD22_TyJ_0350 | 211 | 738 | 244 | 193 |
| BXD27_TyJ_0470 | 218 | 833 | 371 | 243 |
| BXD33_TyJ_0451 | 110 | 288 | 124 | 135 |
| BXD39_TyJ_0478 | 133 | 535 | 0 | 134 |
| BXD40_TyJ_0358 | 94 | 299 | 141 | 100 |
| BXD42_TyJ_0471 | 116 | 490 | 0 | 173 |
| BXD43_RwwJ_0489 | 71 | 391 | 0 | 0 |
| BXD44_RwwJ_0363 | 91 | 385 | 0 | 140 |
| BXD48_RwwJ_0381 | 107 | 404 | 121 | 125 |
| BXD55_RwwJ_0349 | 94 | 242 | 99 | 129 |

| | | | | |
|---|---|---|---|---|
| BXD5_TyJ_0389 | 198 | 847 | 425 | 221 |
| BXD60_RwwJ_0465 | 96 | 561 | 0 | 0 |
| BXD61_RwwJ_0461 | 81 | 356 | 0 | 98 |
| BXD62_RwwJ_0377 | 91 | 331 | 0 | 80 |
| BXD63_RwwJ_0399 | 67 | 213 | 167 | 98 |
| BXD65_RwwJ_0386 | 76 | 289 | 119 | 80 |
| BXD66_RwwJ_0442 | 107 | 435 | 0 | 91 |
| BXD67_RwwJ_0460 | 67 | 231 | 0 | 63 |
| BXD68_RwwJ_0462 | 87 | 332 | 502 | 0 |
| BXD69_RwwJ_0400 | 76 | 258 | 97 | 97 |
| BXD70_RwwJ_0378 | 65 | 272 | 90 | 92 |
| BXD71_RwwJ_0433 | 50 | 185 | 130 | 73 |
| BXD74_RwwJ_0446 | 59 | 249 | 0 | 90 |
| BXD75_RwwJ_0423 | 76 | 332 | 0 | 109 |
| BXD77_RwwJ_0379 | 107 | 326 | 0 | 115 |
| BXD79_RwwJ_0444 | 82 | 412 | 0 | 0 |
| BXD81_RwwJ_0487 | 43 | 163 | 97 | 59 |
| BXD83_RwwJ_0463 | 55 | 413 | 0 | 0 |
| BXD84_RwwJ_0435 | 41 | 230 | 83 | 64 |
| BXD85_RwwJ_0429 | 84 | 251 | 118 | 85 |
| BXD99_RwwJ_0436 | 66 | 286 | 104 | 80 |

38. Lynch, M. Evolution of the mutation rate. *Trends Genet.* **26**, 345–352.
